# A phosphorylation code regulates the multi-functional protein RETINOBLASTOMA-RELATED1 in *Arabidopsis thaliana*

**DOI:** 10.1101/2021.12.20.472892

**Authors:** Jorge Zamora-Zaragoza, Katinka Klap, Jaheli Sánchez-Pérez, Jean-Philippe Vielle-Calzada, Ben Scheres

## Abstract

The RETINOBLASTOMA-RELATED (RBR) proteins play a central role coordinating cell division, cell differentiation and cell survival within an environmental and developmental context. These roles reflect RBR ability to engage in multiple protein-protein interactions (PPIs), which are regulated by multi-site phosphorylation. However the functional outcomes of RBR phosphorylation in multicellular organisms remain largely unexplored. Here we test the hypothesis that phosphorylation allows diversification of RBR functions in multicellular context. Using a representative collection of transgenic loss- and gain of function point mutations in RBR phosphosites, we analysed their complementation capacity in *Arabidopsis thaliana* root meristems. While the number of mutated residues often correlated to the phenotypic strength of RBR phosphovariants, phospho-sites contributed differentially to distinct phenotypes. For example, the pocket-domain has a greater influence on meristematic cell proliferation, whereas the C-terminal region associates to stem cell maintenance. We found combinatorial effects between the T406 phopspho-site with others in different protein domains. Moreover, a phospho-mimetic and a phospho-defective variant, both promoting cell death, indicate that RBR controls similar cell fate choices by distinct mechanisms. Thus, additivity and specificity of RBR phospho-sites fine tune RBR activity across its multiple roles. Interestingly, a mutation disrupting RBR interactions with the LXCXE motif suppresses dominant phospho-defective RBR phenotypes. By probing protein-protein interactions of RBR variants, we found that LXCXE-containing members of the DREAM complex constitute an important component of phosphorylation-regulated RBR function, but also that RBR participates in stress or environmental responses independently of its phosphorylation state. We conclude that developmental-related, but not stress- or environmental-related functions of RBR are defined and separable by a combinatorial phosphorylation code.

## Introduction

Multicellular organisms coordinate cell division and differentiation in space and time to ensure proper development (Gutierrez, 2005; Sablowski and Carnier Dornelas, 2014). When the environment is variable, external cues need to be incorporated in this coordination process. Orchestration of developmental programs and environmental responses becomes particularly challenging in sessile species like plants. The multifunctional protein RETINOBLASTOMA-RELATED1 (RBR) of Arabidopsis, a homolog of the human Retinoblastoma (RB) susceptibility gene product (pRb), acts as an integrator of environmental cues and internal programs into cell fate decisions (Gutierrez, 2005; Harashima and Sugimoto, 2016).

RBR belongs to the pocket-protein family, which function as protein interaction platforms that bring together multiple transcriptional and chromatin regulators, thus controlling genetic programs (Dick and Rubin, 2013; Gutzat et al., 2012). For example, RBR controls cell division by interacting with and inhibiting activation of the S-phase program by E2F-DP heterodimeric transcription factors. Stable repression of cell cycle genes leads to a quiescent state achieved by the DREAM complex (named after its constituents <u>D</u>P-<u>E</u>2F-<u>R</u>BR and the Multivuvla B complex, <u>M</u>uvB), which regulates chromatin structure and DNA methylation (Kobayashi et al., 2015; Ning et al., 2020). RBR-mediated repression is alleviated by phosphorylation, primarily by CYCLIN-DEPENDENT KINASES (CDK). CDKA associates with D-type CYCLINS (CYCD) which target CDKA-CYCD phosphorylation activity to RBR through the CYCD LXCXE motif, thereby releasing E2F repression. RBR also controls formative divisions through similar mechanisms (Cruz-Ramírez et al., 2012; Han et al., 2018; Matos et al., 2014; Weimer et al., 2018) to couple cell division and fate decisions.

The involvement of RBR, and distinct CYC-CDKs and CDK inhibitors (CKI) in both developmental and stress-related processes (Biedermann et al., 2017; Gutierrez, 2005; Horvath et al., 2017; Perilli et al., 2013; Sablowski and Carnier Dornelas, 2014; Wang et al., 2014a; Weimer et al., 2016; Wen et al., 2013; Yi et al., 2014; Zhao et al., 2017), some of which occur simultaneously, points to a central, as yet unspecified role for RBR phosphorylation in the integration of signaling inputs to orchestrate coordinated cell behavior.

Both human pRb and Arabidopsis RBR contain 16 putative CDK phosphorylation sites, most located in the inter-domain regions. Crystal structures of pRb fragments demonstrate that specific phosphorylated residues induce discrete structural changes that promote different intramolecular interactions to either prevent or compete with intermolecular interactions (Burke et al., 2010, 2012). Biochemical characterization of the effect of specific phosphorylation residues on the interaction with E2Fs and with the LXCXE motif indicates that the phospho-sites contribute differentially to regulate pRb-protein interactions (Burke et al., 2010, 2014; Rubin et al., 2005). These observations led researchers to speculate that a ‘phosphorylation-code’ exists, whereby distinct phosphorylation events generate unique structural changes to influence pRb binding properties and functions (Munro et al., 2012; Rubin, 2013). Although attractive, the phosphorylation code hypothesis requires experimental evidence, particularly in plants.

Here, we took a systematic approach to study the biological relevance of RBR phosphorylation. Using a large collection of transgenic loss- and gain of function point mutations in RBR phosphosites, we set out to disentangle RBR roles by specific phosphorylation combinations, and to address whether a phosphorylation code fine-tunes RBR activity. We found that, whereas phosphorylation within the N-domain of RBR gives less prominent effects in general, phosphorylation within the pocket-domain has a greater influence on meristem cell proliferation, and the C-terminal region markedly associates with the stem cell maintenance activity of RBR. Surprisingly, specific combinations of phosoho-defective mutations can lead to hyper-active variants of RBR that promote cell death while restraining proliferation; and the contribution of a phospho-site to the function of RBR varies according the the phosphorylation state of other sites. Finally, we show strong dominant effects of non-phosphorylatable RBR variants and that these can be suppressed by interfering with their ability to bind LXCXE motif-containing proteins like the DREAM complex members of the TCX5/6/7 clade. Our findings provide new insights on the conserved mechanisms underlying RBR function, uncovering the combinatorial nature of RBR phosphorylation-dependent control of cell division, differentiation and survival, while pointing to a phosphorylation-independent role in stress and environmental responses.

## Results

### A system to study phospho-variants by circumventing early lethality

The substantial knowledge on plant RETINOBLASTOMA-RELATED (RBR) proteins derives from expression studies, null or hypomorphic alleles, and up- or down-regulation of the gene (Ach et al., 1997; Borghi et al., 2010; Chen et al., 2011; Cruz-Ramírez et al., 2013; Ebel et al., 2004; Grafi et al., 1996; Gutzat et al., 2011; Perilli et al., 2013; Wachsman et al., 2011; Wildwater et al., 2005; Xie et al., 1996). However, the functional outcome of RBR phosphorylation remains largely unexplored, in spite of being assumed to be a major regulatory mechanism of RBR activity. We approached this subject by constructing a representative collection of transgenic RBR phospho-variants comprising all putative CDK-phosphorylation sites (Desvoyes and Gutierrez, 2020; Desvoyes et al., 2014) (Fig. 1).

**Figure 1.**
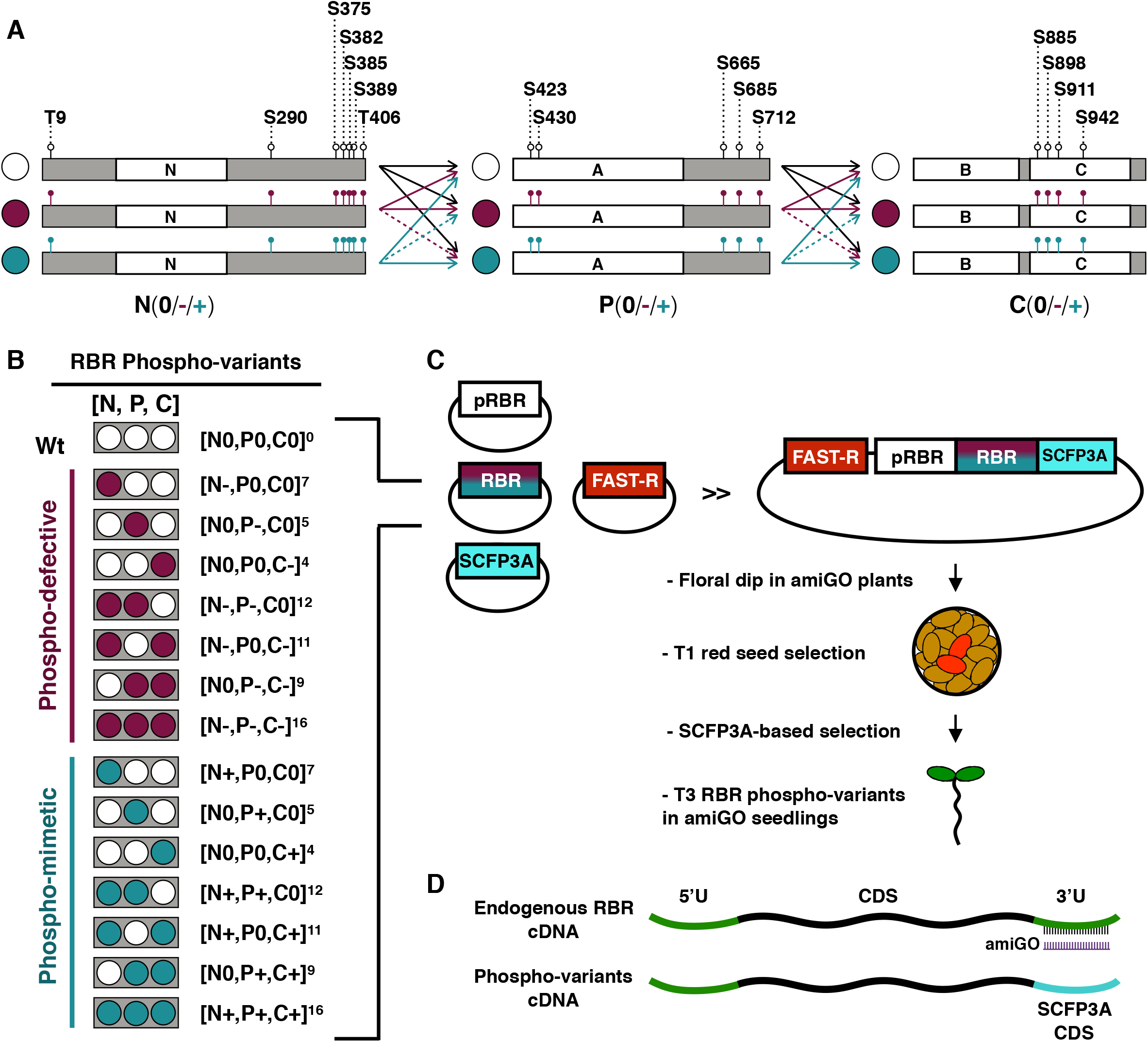
A system to study phospho-variants by circumventing early lethality. A) We cloned three combinable fragments (N, P, and C) comprising the full length RBR cDNA into the level −1 vector of the Golden Gate MoClo toolbox (Engler et al., 2014). White boxes represent RBR protein domains (**N**-terminal, **A**- and **B**-pocket subdomains, and **C**-terminal) as predicted by pfam server (https://pfam.xfam.org/), and gray boxes represent unstructured protein regions. Note that the P module contains all phospho-sites within the **<u>P</u>**ocket domain, but due to cloning convenience, the C-module encodes the B-pocket sub-domain. Each level −1 module encodes a subset of phosphorylation sites indicated by empty or colored lollipops, and the corresponding the amino acid residue (S/T) position. Since all phospho-sites within a module are in the same state, we use colored circles and the signs “0”, and “+” to denote the module state: white/0 for phosphorylatable (Ser/Thr), dark red/− for phospho-defective (Ala), and teal/+ for phosphomimetic (Asp or Glu). All codon changes are listed in Table S1. All combinations are possible (arrows), but we avoided phospho-defective with phosphomimetic combinations (dotted arrows). B) Color code and text nomenclature of phospho-variants. C) Generation and selection of transgenic RBR phospho-variants plants. Modules were assembled into the level 0 vector to generate full length RBR phospho-variants CDS, subsequently assembled with RBR promotor, the CDS of SCFP3A fluorescent tag, and a terminator (not illustrated) into Level 1 constructs. Level 2 constructs containing the FAST-R selection cassette and the corresponding phosphovariant were transformed in 35::amiGO-RBR plants (amiGO). T1 seedlings pre-selected by the red seed coat were selected for the best SCFP3A intensity and taken to T3 generation for complementation analysis. D) amiGO selectively down-regulates endogenous RBR transcripts by targeting the 3’-UTR, which is absent in the transgenic RBR:SCFP3A CDS. See Figure S1 for the full list of modules and phosphovariants generated.

Tests of all possible phosphorylation states on 16 sites would entail the construction of 3^16^ (~ 43 million) variants, so we simplified the analysis by taking a domain approach. Briefly, the coding sequence of RBR was split into three combinable modules named as “N”, “P”, and “C” (after the **<u>N</u>**-terminal, AB-**<u>P</u>**ocket, and **<u>C</u>**-terminal protein domains), each bearing a subset of phosphosites in the one of three states: phosphorylatable (wild-type), phospho-defective, and phosphomimetic, the later resembling constitutive phosphorylation (Antonucci et al., 2014; Chen et al., 2017; Dissmeyer and Schnittger, 2011; Sanidas et al., 2019; Wang et al., 2014b). These are depicted by “0”, “-” and “+” signs, respectively (or by a colored circles code in figures, Fig 1A,B; S1A). We refer to each RBR phospho-variant as the specific combination of modules, with a superscript indicating the total number of mutated sites. For example, [N0,P0,C0]^0^ refers to the fully phosphorylatable RBR, while [N0,P-,C0]^5^ and [N0,P+,C0]^5^ respectively denote phospho-defective and phosho-mimetic versions of the five phosphor-sites in central module only (Fig 1B). We refrained from combining phospho-defective with phospho-mimetic modules and assembled all other possible variants with the native RBR promoter and a SCFP3A C-terminal tag to select comparable expression levels of transgenic plants (Fig. 1B,C, Fig S1B). All RBR phosphovariants were transformed into plants homozygous for the amiGO-RBR genetic construct (hereafter, amiGO; Fig 1C,D), an artificial microRNA driven by the 35S promoter that selectively down-regulates endogenous RBR transcripts only after the gametophyte stage and completion of early embryogenesis (Cruz-Ramírez et al., 2013). This late reduction in RBR levels bypasses is requirement in early developmental stages and enhances both cell proliferation and death similar to a true null *rbr* clone (Wachsman et al., 2011). Through analysis of the complementation capacity of all viable homozygous transgenic variants at stages when the amiGO phenotypes were fully penetrant (Fig S2), we could assess the effect of site-specific mutant combinations in RBR.

### C-region phosphorylation inhibits RBR-mediated restriction of stem cell (SC) division

Since downregulation of RBR leads to supernumerary QC and SC divisions (Cruz-Ramírez et al., 2013; Wildwater et al., 2005), we first asked whether stem cell niche (SCN) proliferation is affected by specific RBR-phosphorylation events. Aberrant division planes hinder lineage identification in the absence of markers, so we quantified the pooled number of QC, cortex and endodermis initials (CEI), and columella stem cells (CSC) to explore the effect of the phospho-variants in SCN maintenance.

All but two unviable phospho-defective variants (indicated as ⌧ in Fig. 2A) complemented the SCN overproliferation phenotype induced by the amiGO at a level at least equal to the complementation by wild type RBR (Figure 2A,B), consistent with dephosphorylated RBR acting as the repressor of SCN activity. Among these variants, the combination of phospho-defective residues in N and P domains [N-,P-,C0]^12^ showed the higher level of repression indicating that phosporylations in these domains have an additive effect. Thus dephosphorylation is essential for the role of RBR in normal maintenance of the SCN, and single-domain dephosphorylation is insufficient for maximal RBR-mediated repression.

**Figure 2.**
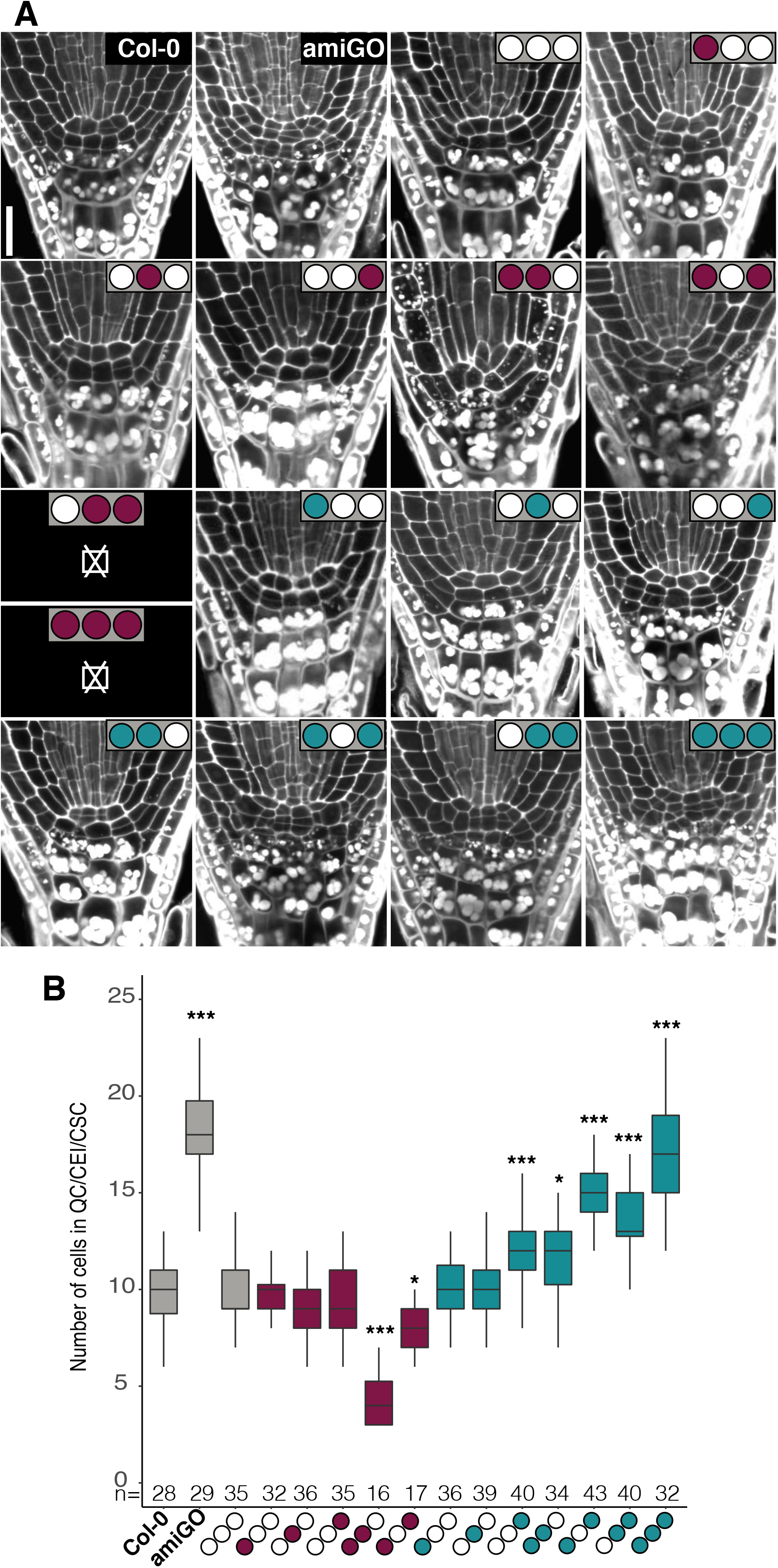
C-region phosphorylation inhibits RBR-mediated restriction of stem cell (SC) division. A) Representative confocal images of modified pseudo-Schiff propidium iodide (mPS-PI) stained root tips of the indicated genotypes. Sub-panels marked with ⌧ correspond to lethal genotypes. B) Box plot of pooled QC cells, CEI, and CSC number excluding cells with evident starch granules accumulation. Data from two biological replicates presented as median and interquartile range, n denotes total number of scored roots. Dunett’s test against Col-0, ***p < 0.001, *p < 0.05. Scale bar, 20 μM.

Consistent with a role for dephosphorylated RBR in repressing SCN activity, several variants with a phospho-mimetic module failed to suppress SCN overproliferation. Overproliferation never exceeded that seen in the amiGO background, but increased with the number of domains containing phospho-mimetic residues. Thus, phosphorylation in more than one RBR domain is needed to relieve the repression of SCN divisions. However, [N0,P0,C+]^4^ revealed incomplete repression of SCN activity similar to [N+,P+,C0]^12^, despite having fewer phospho-mimetic residues, indicating that phosphorylation is not simply additive and that sites in the C domain have a greater influence on SCN regulation than those in the N and P domains. Taken together, a range of phospo-defective and phospho-mimetic mutant combinations reveals additive but differential contributions to the regulation of SCN activity by RBR phospho-sites in all three protein domains.

### Meristem size maintenance depends most strongly on Pocket domain phosphorylation

Similar to their effect on SCN activity, down-and up-regulation of RBR have opposite effects on root meristem size (Perilli et al., 2013), reflecting control of cell division. To elucidate whether cell division activity also depends on a ‘phosphorylation code’ in the root meristem, we measured the effects of the phospho-site variants on the size of the transit amplifying cell pool in the meristem. As expected, amiGO meristems were slightly longer and contained more cells than Col-0, which could be fully restored using the Wt RBR version [N0,P0,C0]^0^ (Fig 3A, 3B).

**Figure 3.**
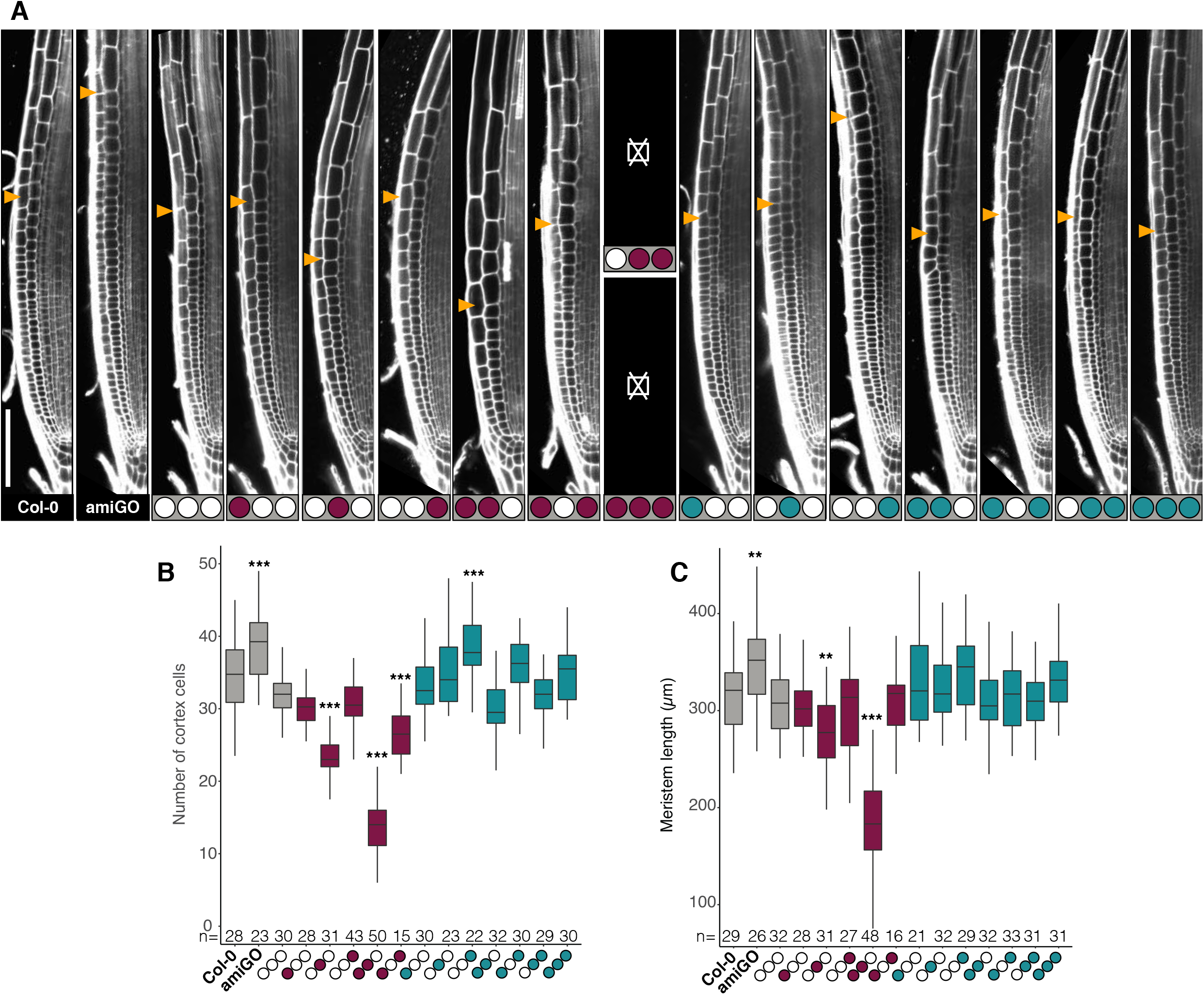
Meristem size maintenance depends most strongly on Pocket domain phosphorylation. A) Representative confocal images of root meristems of the indicated genotypes; yellow arrowheads mark the end of the meristem proliferation zone. Sub-panels marked with ⌧ correspond to lethal genotypes. B,C) Box plots of meristem proliferation and size quantified as the number of cortex cells B) and length C) from QC to the first rapidly elongating cortex cell. Data from two biological replicates presented as median and interquartile range, n denotes total number of scored roots. Dunett’s test against Col-0, ***p < 0.001, **p < 0.01. Scale bar, 100 μM.

With exception of [N0,P0,C+]^4^, which contained more cells but the meristem length was not significantly different than the Wt, phospho-mimetic variants were all able to complement the amiGO meristem phenotype, indicating that almost any additional activity of RBR mitigates the slight meristem size increase observed in the amiGO root meristem. Unlike observed for the SCN proliferation phenotype, no other variant containing the C+ module exhibited significant changes (Fig 3A, 3B). Conversely all phospho-defective variants reduced amiGO-induced overproliferation in the meristem (Fig 3B). However, in this case the phospho-defective Pocket domain alone in [N0,P-,C0]^5^ was sufficient to over-complement the amiGO mutants, exhibiting shorter meristems than the Wt, without significant effect on the SCN (Fig 2B, 3B,C). Our data indicate distinct effects for RBR phosphorylation sites in control over SCN and meristem proliferation, with a larger role for the C-region phosphorylation in the SCN, and for the Pocket domain in the transit amplifying cells of the meristem.

Interestingly, despite presenting more cells (Fig 3B), [N0,P0,C+]^4^ did not increase meristem length (Fig 3C), indicating that a compensatory mechanism maintains meristem size. Similar compensation effects were seen for [N-,P0,C-]^11^, where less cells did not lead to a difference in meristem length compared to Col-0. Compensatory mechanisms did not sustain meristem size whenever the phospho-defective P module was present. Thus, phosphorylation of the Pocket domain is particularly important to maintain meristem size, consistent with a site-specific component in the phosphorylation code.

### Suppression of cell death is rescued by all but two distinct RBR phosho-variants

Spontaneous cell death in the root tip constitutes a hallmark of reduced RBR activity (Cruz-Ramírez et al., 2013; Wildwater et al., 2005), likely due to the inability to cope with intrinsic DNA damage (Biedermann et al., 2017; Horvath et al., 2017). We examined the protective role of RBR phosphorylation variants using propidium iodide staining (PI), which permeates only dead cells. As expected, all amiGO roots presented dead cells, while Col-0 and the vast majority of the phospho-variants had around 25% or less root tips with dead cells. Two phospho-variants reached a comparable cell death frequency to amiGO seedlings (Fig 4A, 4B). The full phosphomimetic variant [N+,P+,C+]^16^ fits the paradigm of hyper-phosphorylated RBR being inactive. In agreement with this, [N+,P+,C+]^16^ also presented overproliferation of SCN and meristematic cells (Figs. 2 and 3A,B), supporting the supposition that phospho-mimetic mutations inactivate RBR. However, the phospho-defective variant [N-,P0,C-]^11^ presented a striking outcome for a RBR isoform presumed to be active although not able to be phosphorylated in N nor C terminal domains. [N-,P0,C-]^11^ over-complemented the amiGO cell proliferation phenotypes (Figs 2 and 3A,B) but failed to promote cell survival, in contrast with [N0,P-,C0]^5^ and [N-,P-,C0]^12^, that also over-complemented cell proliferation but fully restored the cell death phenotype (Fig 2B, 4A,B). Additionally, some phospho-mimetics that failed to restrain SCN proliferation still suppressed cell death. Thus, cell proliferation is always promoted by RBR phosphorylation to a greater or lesser extent according to specific phospho-sites, but cell death emerges either upon constitutive RBR hyper-phosphorylation or with a specific combination of un-phosphorylated sites, implying two different mechanisms for RBR-promoted cell survival.

**Figure 4.**
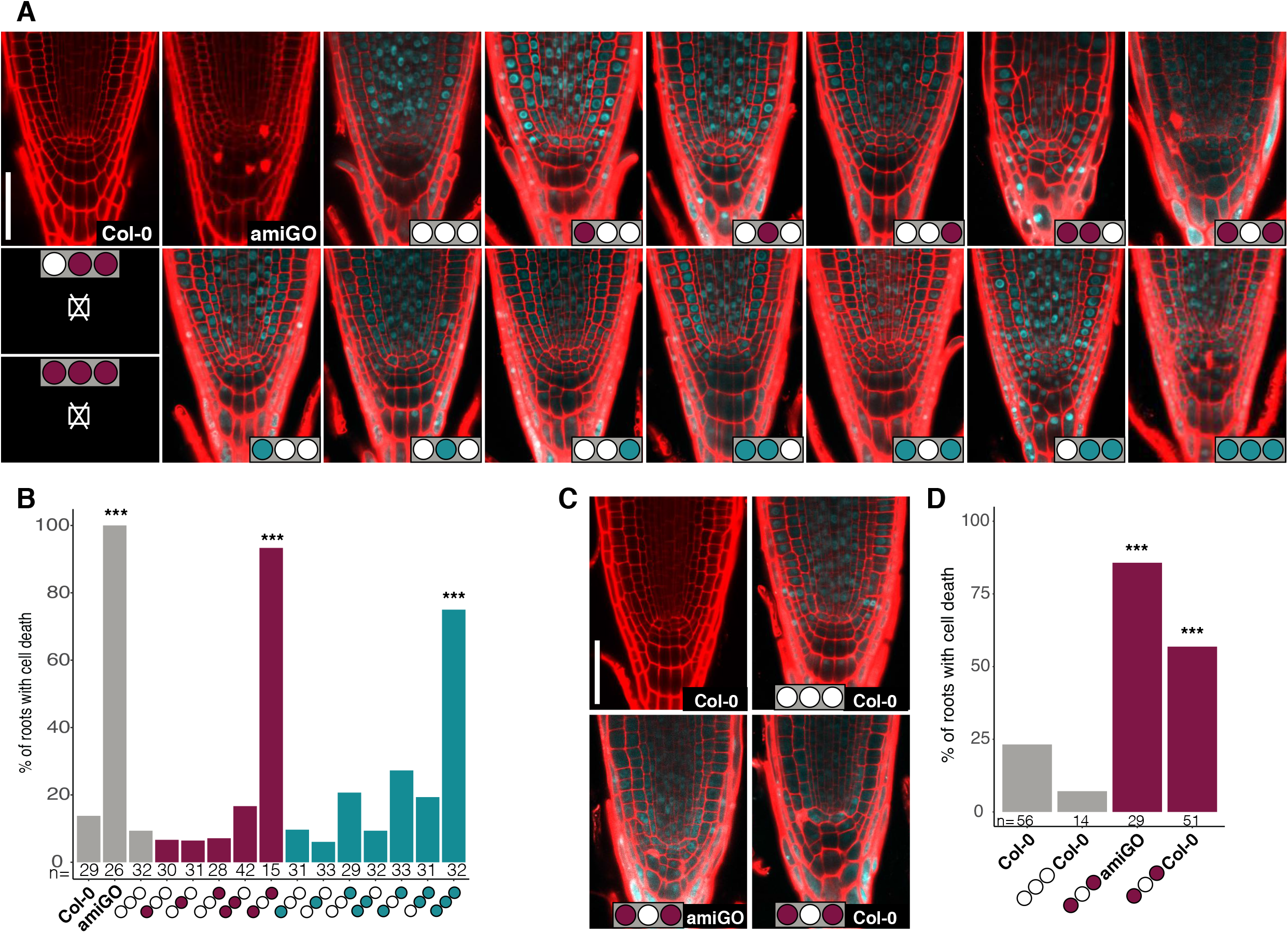
Suppression of cell death is rescued by all but two distinct RBR phosho variants. A and C) Representative confocal images of PI-stained root tips of the indicated genotypes. Subpanels marked with ⌧ correspond to lethal genotypes. In C) the genetic background of phosphovariants is indicated in black text boxes. B and D) Bar graphs from A) and C), respectively, showing percentage of root tips with dead cells. Data from two biological replicates presented as means, n denotes total number of scored roots. Fisher’s test against Col-0, ***p < 0.001. Scale bars, 50 μM.

Since phospho-defective [N-,P0,C-]^11^ efficiently restrains cell division, we asked whether the cell death phenotype is caused by the activity of RBR, or results from an impaired protective function. We out-crossed the amiGO background to assess the effect of [N-,P0,C-]^11^ and [N0,P0,C0]^0^ in the presence of endogenous RBR. While 4 copies (endogenous and transgenic) of wild-type RBR conferred a protective effect, more than 50% of the [N-,P0,C-]^11^ roots still displayed dead cells in the Col-0 genetic background (Fig 4 C,D). However, in the amiGO background the frequency increased to more than 80% (Fig 4C,D), indicating that [N-,P0,C-]^11^ is an RBR active isoform triggering the cell death program, possibly counteracted by the endogenous RBR.

### Combining full domain with single-site phospho-site variants indicates a combinatorial phosphorylation code

The distinct contributions of phospho-sites in different RBR protein domains to cell division phenotypes (C-region sites to SNC activity and Pocket domain sites to meristem size) contrasts with the more equal contribution of sites to the cell death effect (Fig 4, compare [N-,P0,C-]^11^ to [N-,P0,C0]^7^ and [N0,P0,C-]^4^). To explore potential effects of a single specific phospho-site in one module to RBR phenotypes when combined with defective sites in other modules, we generated a new “N” phospho-module containing a single phospho-defective site on position T406, thus named as “406-” (Figs. 5A, S1A). All combinations of the 406-module with the WT and phospho-defective P and C modules were analysed for phenotype —except for [406-,P-,C-]^10^ that was inviable.

**Figure 5.**
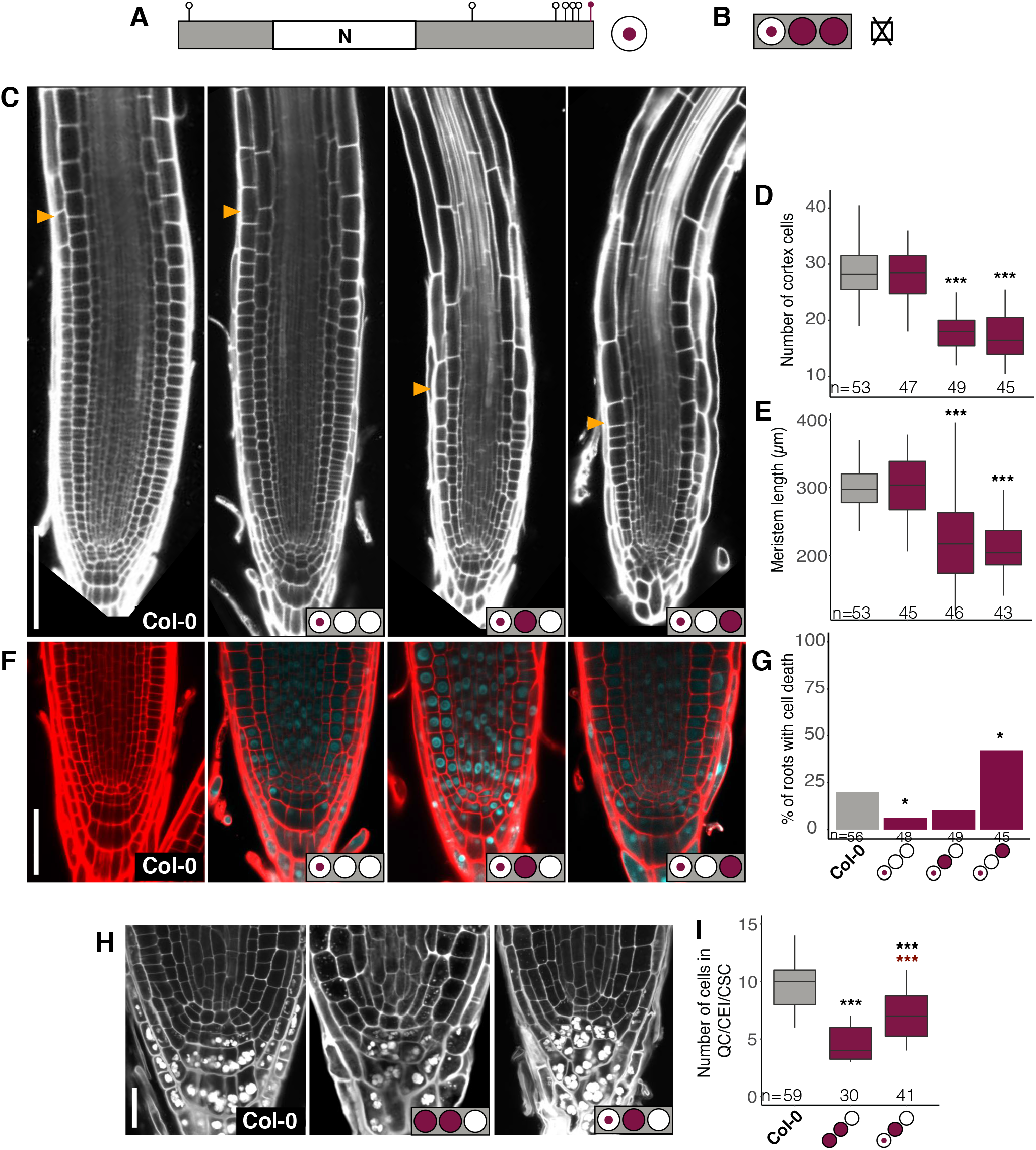
Combining full domain with single-site phospho-site variants indicates a combinatorial phosphorylation code. A) Schematic representation of the phospho-defective 406 module and its colored circle code. B) The ⌧ symbol indicates that the [406-,P-,C-]^10^ phospho-variant is lethal. C, F, H) Representative confocal images of PI-C,F) or mPS-PI-H) stained root tips of the indicated genotypes; yellow arrowheads mark the end of the meristem proliferation zone. D and E) Box plots from C) of meristem proliferation and size quantified as the number of cortex cells D) and length E) from the QC to the first rapidly elongating cortex cell. L) Bar graph from F) showing the percentage of root tips with dead cells. I) Box plot from H) quantifying the pooled QC cells, CEI, and CSC number excluding cells with evident starch granules accumulation. Data from two biological replicates presented as median and interquartile range D,E,I) or as means G); n denotes total number of scored roots. Fisher’s test G) or Dunett’s test against Col-0 D,E,I) and against [N-,P-,C0]^12^ I), ***p < 0.001, black asterisks indicate significant differenced against Col-0; red asterisks in I) against [N-,P-,C0]^12^. Shared labels in ‘x’ axis for D),E),G). Scale bars, 100 μM in (C), 50 μM in (F), 20 μM in (H).

Similar to the fully phospho-defective N-domain, meristem size was not affected by 406-alone, but was reduced by one third in [406-,P-,C0]^6^ (Fig. 5C-E). Since this effect was milder than in [N-,P-,C0]^12^ (Fig 3), we conclude that T406 has an additive effect to the strong influence of the Pocket domain phosphorylation on meristem size. Notably, [406-,P0,C-]^5^ showed an equally strong effect (Fig 5C-E), and even more severe than [N-,P0,C-]^11^ (Fig 3) suggesting that, when combined with those in the C-region, not all phospho-sites in the N-domain are additive with respect to the repressive function of RBR in meristem size maintenance.

Unlike [406-,P-,C0]^6^, [406-,P0,C-]^5^ displayed increased cell death (Fig 5L), but to a lesser extent than its high order counterpart [N-,P0,C-]^11^ (Fig 4). Since [N0,P0,C-]^4^ and [406-,P0,C0]^1^ showed full or even enhanced cell survival in the latter case (Fig 5 F,G; see Fig 4 A,B, for [N0,P0,C-]^4^), we conclude that none of the phospho-sites by their own, but the combination of dephosphorylated sites in the N and C regions trigger cell death, and that individual sites in the N domain exhibit an additive effect on the phenotype penetrance.

In turn, [NT406-,P-,C0]^6^ restricted SC divisions but to a lesser extent than [N-,P-,C-0]^12^ (Fig 5 H,I), suggesting the additive effect of N and Pocket domains phosphorylation on RBR activity, while the full complementation conveyed by [N-,P0,C0]^7^ and [N0,P-,C0]^5^ (Fig 2), indicates that combinatorial dephosphorylation of the RBR N and Pocket domains restricts SCN activity. Unfortunately, we could not assess the effect of [406-,P0,C-]^5^ on SC divisions due to limited seed availability. Altogether, the phospho-defective 406 residue enhanced the activity of the P- and C-contained phospho-sites to restrict cell division (to even a greater extent than the N-module when combined with C-), and triggered cell death activation only in combination with the C-terminal phospho-defective module, indicating that the phenotypic effect of an individual phospho-site depends on the phosphorylation status of the remaining ones.

### Fertility and embryogenesis are compromised in highly substituted RBR phosphodefective variants

The limited seed production of [N-T406,P0,C-]^5^ was also observed in [N-,P-,C0]^12^ and [N-,P0,C-]^11^. Moreover, the few [N0,P-,C-]^9^ transformants we obtained that showed detectable sCFP fluorescence resulted in fully sterile plants (Fig S3 A-D), highlighting the importance of phosphorylation throughout the P and C regions to sustain plant reproduction. Lack of fertilization in more than 80% of [N-,P-,C0]^12^ and [N-,P0,C-]^11^ ovules, plus a smaller fraction of aborted seed added up to nearly 90% of sterility, regardless of the genetic background (amiGO or Col-0; Fig S4A). Consistently, both male and female reproductive tissues displayed cytological defects (Fig S4B,C). Thus, defective reproductive development results not only from reduced RBR activity as previously reported (Ebel et al., 2004; Zhao et al., 2017), but also from hyper-active isoforms, indicating that RBR is regulated by phosphorylation during gametophyte development.

Since agrobacterium-mediated transformation occurs specifically in the female reproductive tissues (Desfeux et al., 2000), gametophytic defects may account for the lack of recovery of transgenic seedlings expressing [N-,P-,C-]^16^ or [406-,P-,C-]^10^ phospho-variants. But even if transformed ovules are fertilized, embryo lethality can also occur since RBR regulates embryonic genetic programs (Gutzat et al., 2011). To explore this possibility, we used the red fluorescent seed coat selection marker (see Fig 1C) to select [N-,P-,C-]^16^ primary transformants in both amiGO and Col-0 backgrounds, and recovered all embryos from non-germinated seeds. A small fraction of embryos (~3.5%, n=318) was arrested at heart- to torpedo stages and showed enlarged cells regardless the presence of endogenous RBR (Fig. 6 A, Movie S1). Some arrested embryos presented residual or absent radicles (Fig S3F,G), single or uneven cotyledons (Fig S3H,J), and signs of early differentiation like root hairs (Fig S3I). Conversely, we did not find any of these features in non-germinated seeds of Col-0 nor in primary transformants of a viable phospho-defective RBR variant (Fig. 6A, S3K). Considering the phenotypic similarities in arrested embryos of [N0,P-,C-]^9^ transformants (Fig S3E, MovieS2), our results indicate that a dominant effect of phospho-defective mutants (particularly in the P and C regions) blocks embryonic development. Altogether, defective reproduction and early developmental arrest underlie the viability loss of highly substituted phospho-defective variants, leading us to conclude that RBR multi-phosphorylation, particularly on the Pocket domain and C-terminus, is essential for plant survival.

**Figure 6.**
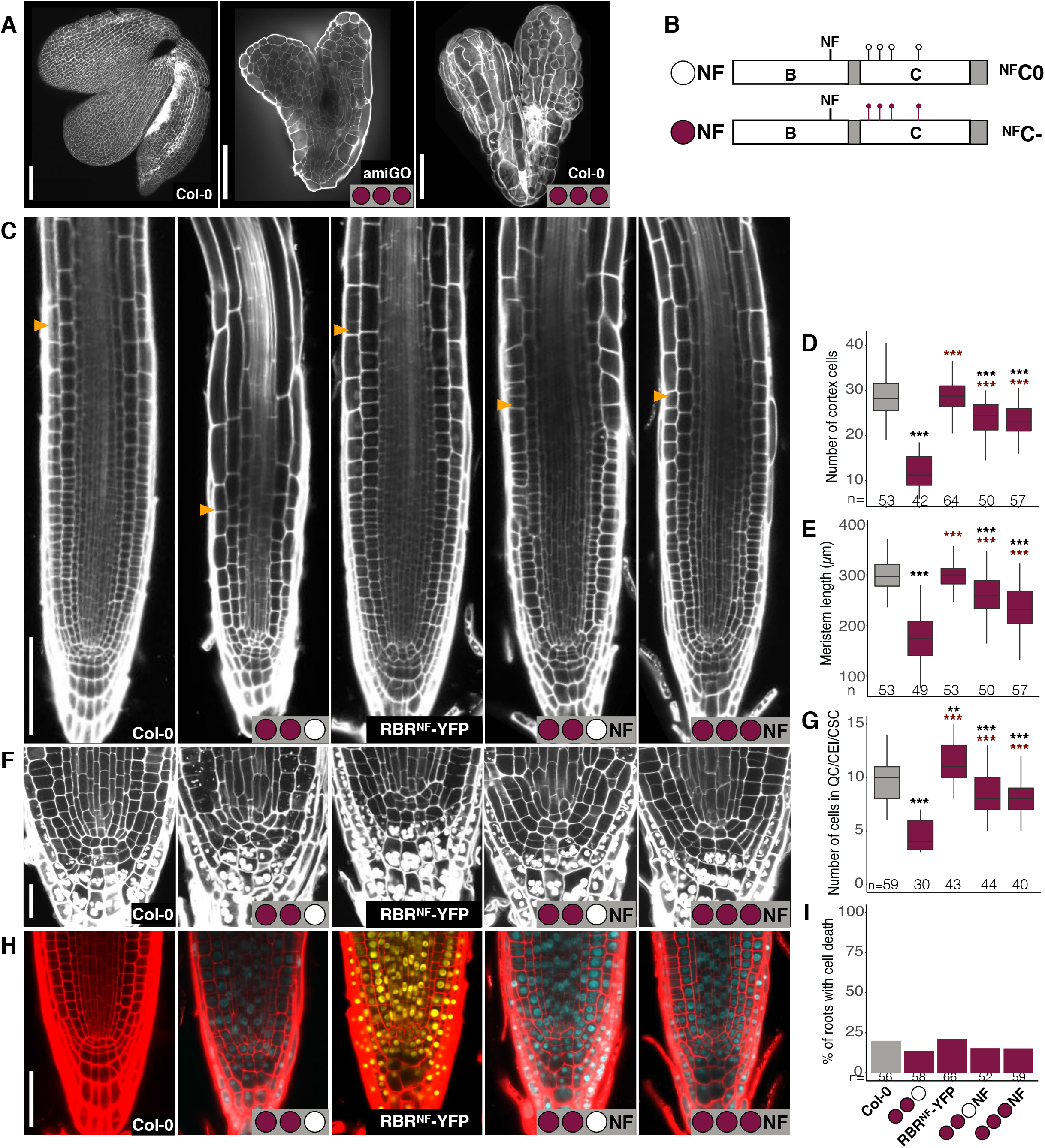
A point mutation in the B-pocket sub-domain rescues highly substituted phospho-defective mutants. A) Confocal images of mPS-PI stained embryos from non-germinated seeds 4 days after sowing (das) stratified for 4 days of Col-0 and primary transformants of [N-,P-,C-]^16^ in the genetic backgrounds amiGO and Col-0. B) Schematic representation of the ^NF^C0 and ^NF^C-modules and its colored circle code. C,F,H) Representative confocal images of PI-C,H) or mPS-PI-F) stained root tips of the indicated genotypes; yellow arrowheads mark the end of the meristem proliferation zone. D,E) Box plots from C) of meristem proliferation and size quantified as the number of cortex cells D) and length E) from the QC to the first rapidly elongating cortex cell. G) Box plot from F) quantifying the pooled QC cells, CEI, and CSC number excluding cells with evident starch granules accumulation. I) Bar graph from H) showing the percentage of root tips with dead cells. Data from two biological replicates presented as median and interquartile range D,E,G), or as means I); n denotes total number of scored roots. Fisher’s test I) or Dunett’s test against Col-0 and against [N-,P-,C0]^12^ D,E,G), ***p < 0.001, **p<0.01, black asterisks indicate significant difference against Col-0; red asterisks, against [N-,P-,C0]^12^. Col-0 and [N-,P-,C0]^12^ values are the same as in Figure 5 as experiments were performed in parallel sharing these controls. Shared labels in ‘x’ axis for D),E),G), I). Scale bars, 100 μM in (A,C), 20 μM in (F), 50 μM in (H).

### A point mutation in the B-pocket sub-domain rescues highly substituted phosphodefective mutants

If RBR phosphorylation disrupts its protein interactions, the dominant phenotypes of highly substituted phospho-defective RBR variants might reflect more stable protein interactions. To investigate this hypothesis, we introduced the point mutation N849F (human N757F, mouse N750F —hereafter NF), which disrupts interactions with LXCXE motif-containing proteins in plants and animals (Bourgo et al., 2011; Chen and Wang, 2000; Cruz-Ramírez et al., 2013), into two “C” modules to generate two new phospho-defective RBR alleles: [N-,P-,^NF^C0]^12^ and [N-,P-,^NF^C-]^16^ (Fig 6A, S1). Strikingly, we recovered viable plants and homozygous lines, even for the fully phospho-defective variant.

To investigate the suppressive effect of the NF mutation, we compared the phospho-defective NF variants alongside pRBR::RBR^NF^:vYFP (hereafter RBR^NF^), with both Col-0 and [N-,P-,C0]^12^. As reported previously (Cruz-Ramírez et al., 2013; Zhou et al., 2019), the RBR^NF^ allele showed a slight overproliferation of the SCN (Fig 6 F,G). The NF mutation partially restored the meristem size phenotypes of the over-complementing [N-,P-,C0]^12^ variant (Fig 3 vs Fig 6 C-E); similarly, SCN differentiation was also partially rescued (Fig 2 vs Fig 6 F,G). Moreover, the fully phospho-defective [N-,P-,^NF^C-]^16^ variant showed little, if any, phenotypic variation compared to [N-,P-,^NF^C0]^12^ (Fig 6 C-I). Even non-germinating [N-,P-,^NF^C-]^16^ primary transformants showed more advanced development than [N-,P-,C-]^16^ arrested embryos (Fig 6 A vs S3K). Thus, the partial rescue of highly substituted phospho-defective variants phenotypes by the NF mutation suggests that RBR-LXCXE protein interactions constitute a predominant component of RBR-mediated developmental processes regulated by phosphorylation.

### RBR protein-protein interactions with transcriptional regulators are differentially regulated by phosphorylation

To search for RBR protein interactions that explain our observations, we performed Yeast Two-hybrid (Y2H) screenings of the Arabidopsis pEXP22-TF collection (Pruneda-Paz et al., 2014) using eight highly substituted RBR phospho-variants, RBR^NF^, and wild type RBR as baits. We confirmed 14 interactors with varying binding properties, some of which were previously reported and others were unknown (Fig. S5). Several interactors are involved in responses to stress, four of which (DREB2D, GBF4, NAC090, NAC044) interacted strongly with all RBR phospho-variants, pointing to a phosphorylation-independent role of RBR as an integrator of environmental inputs. Conversely, TFs related to cell proliferation and development (TCX6/7, E2FC, XND1, TCP3) showed weaker or no interaction with most phospho-mimetic variants. Taken together with our phenotypic analysis, these results indicate that RBR protein structure is functional despite the multiple mutations, and that phospho-site substitutions work as expected in both of their defective and mimetic versions. Two interactors revealed special properties. On the one hand, ARIA interacted strongly and exclusively with RBR variants containing an intact N-domain, suggesting that this protein docks closely to, or even on phospho-sites at the N-domain. On the other hand, GRF5 showed strong but variable binding properties to RBR, since it interacted with all RBR variants in at least one replicate, but consistently only with Wt RBR, [N-, AB0, C-], and [N-, AB-, C-] (Fig S5B), suggesting that other factors might be required to stabilize the RBR-GRF5 complex, like the co-activator GRF-INTERACTING FACTOR1/ANGUSTIFOLIA3 (GIF1/AN3).

### Phosphorylation-regulated functions of RBR are largely mediated by the interaction with members of the DREAM complex

We paid special attention to the cysteine-rich proteins TCX6 and TCX7, members of the multimeric DREAM complex, a conserved eukaryotic cell cycle regulator recently described in plants (Kobayashi et al., 2015; Lang et al., 2021; Ning et al., 2020). TCX6/7 displayed decreased affinity for phospho-mimetic RBR variants and no interaction with RBR^NF^. Together with TCX5, TCX6 and TCX7 contain a conserved LXCXE motif responsible for the interaction with RBR that is absent in the remaining TCX family members (Fig S6A-C; (Lang et al., 2021). Since TCX7 expression was undetectable in all tissues analysed (Andersen et al., 2007), and the double, but not the single mutants of *tcx5* and tcx6 exhibit phenotypes in other plant parts (Ning et al., 2020), we analysed the root meristem of the *tcx5/6* double mutant. The *tcx5/6* mutant exhibited increased meristem size, cell proliferation, and cell death (Fig S6D-H), similar to the phenotypes of the amiGO root tips, suggesting that the RBR-TCX5/6 interaction is relevant for RBR function. Thus, we transformed the dominant lethal phospho-defective [N-,P-,C-]^16^ variant of RBR in the *tcx5/6* mutant background. Surprisingly, we recovered 16 independent primary transformants with detectable CFP3A nuclear signal, indicating that the absence of TCX5/6 proteins is enough to circumvent the dominant lethality of such a variant. Nevertheless, only one transformant line generated T2 seed, indicating that the suppression of the [N-,P-,C-]^16^ variant by *tcx5/6* is weaker than in that observed in [N-,P-,^NF^C-]^16^. Accordingly, the phenotypes of the T2 *tcx5/6*;[N-,P-,C-]^16^ line are stronger than those observed for [N-,P-,^NF^C-]^16^(compare Fig S6D with Fig 6), despite the weaker SCFP3A intensity of the former. Altogether, we concluded that the roles of RBR regulated by phosphorylation and mediated by its LXCXE-binding properties, partially depend on the interaction with the DREAM complex members of the TCX5/6/7 clade.

## Discussion

RBR is associated with multiple and complex roles in development, and it has been unclear how a single protein can carry out a wide range of functions through regulated interactions with different protein partners. Here, we have explored the separability of roles for plant RBR phosphorylation, which has emerged as a prominent regulatory mechanism of RBR in the multitude of RB proteins functions described so far. Taken together, our results support the notion that a phosphorylation code fine-tunes RBR activity and function providing the potential for differential regulation in its various roles. While phospho-defective variants often showed dominant effects and over-complementation of ‘amiGO’ plants with reduced RBR levels (Figs. 2,3,4C,D,5, 6A, S3,S4A), the phenotypic strength of phospho-mimetic variants ranged between those observed for wild-type and amiGO (Figs. 2–4), which supports the prevailing conception that an active, unphosphorylated RBR, is inactivated by regulatory phosphorylations. But three additions to this generic idea are to be made: phosphorylation events on RBR (1) are independent of each other, (2) unequally contribute to RBR activity, and (3) disentangle RBR functions (provide evidence for potentially independent regulation of different RBR functions?).

### RBR phospho-sites are independent of each other

We observed an additive effect in the phenotypic strength as the number of mutated phosphosites was increased in RBR variants. Together with the phenotypic differences between full phospho-site variants ([N-,P-,C-]^16^ and [N+,P+,C+]^16^) and all single phospho-module combinations, our findings exclude a nucleation mechanism for RBR hyper-phosphorylation, and demonstrate that phosphorylation events on RBR are independent of each other.

### Uneven contribution of phospho-sites to RBR activity regulation

Unlike [N-,P-,C0]^12^ and [N-,P0,C-]^11^, the less substituted phospho-defective variant [N0,P-,C-]^9^ was lethal. Therefore, the phosphorylatable Pocket domain and C-region, but not the N-region of RBR protein, were sufficient to sustain plant growth and viability despite bearing less phosphosites. This excludes a simple phosphor-counting mechanism as the primary mode of RBR regulation. Moreover, phosphorylation within the Pocket domain and the C-region markedly influenced the proliferative activity of the meristem and SCN, respectively, whereas phosphorylation of the N domain seemed unimportant on its own (Figs 2,3). Accordingly, phosphorylation within the Pocket domain and C-region of pRb regulate E2F and LXCXE motif binding (Burke et al., 2010; Knudsen and Wang, 1997), while the relevance of phosphorylating the N-domain has been shown to emerge in response to stress (Gubern et al., 2016). Future research should unveil the functions of RBR N-domain phosphorylation during plant stress responses.

### RBR functions are separable by phosphorylation

While the phospho-defective variant [N0,P-,C0]^5^ over-complemented root meristem size but not SCN division, all three double phospho-mimetic modules combinations displayed overproliferation of the SCN but not of transit amplifying cells. Notably, several phospho-mimetic variants that failed to restrict SCN activity, complemented the cell death phenotype. On the other hand [N-,P0,C-]^11^ and [N-^T406^,P0,C-]^5^ repressed cell division and frequently displayed dead cells, whereas [N-,P-,C0]^12^ and [N-^T406^,P-,C0]^6^ were blocked meristematic and stem cell division without inducing cell death (Figs 2–4). Contrary to previously observed pleiotropic effects in knock-out or altered expression approaches, our findings revealed the capacity of RBR to regulate independently cell division, differentiation and survival according to its phosphorylation state.

We observed increased cell death in roots of both phospho-mimetic and phospho-defective variants. Similarly, apoptotic stimuli can promote phosphorylation as well as de-phosphorylation of pRb (Leon et al., 2008; Nath et al., 2003); pRb in turn, can either promote or inhibit apoptosis (Antonucci et al., 2014; Goodrich, 2006; Ianari et al., 2010), a fate decision largely mediated by its phosphorylation state (Antonucci et al., 2014; Egger et al., 2016; Gubern et al., 2016; Lee et al., 2018; Leon et al., 2008; Nath et al., 2003). Which particular phosphorylation state is preferred, and what outcome it takes might depend on circumstances. In plants, biotrophic attackers promote RBR hyper-phosphorylation to trigger immunity-related PCD (Wang et al., 2014a), correlating with our fully phospho-mimetic variant. We speculate that rapid immune responses to pathogen attack, sensed and signaled by phospho-relay cascades, prioritize an urgently required activation of PCD to avoid infection spread. On the other hand a proper balance between cell division, differentiation, and developmental PCD could involve a more accurate, finely-tuned mechanism. Probably this includes coordinated action of CDKs and phosphatases, reflected by the combinatorial specificity of our phospho-defective variants triggering cell death and inhibiting cell division (Fig 4, 5L). Thus, hyper-phosphorylation of RBR may well act “quick and dirty” to counteract stresses, while combinatorial phosphorylation entails a timely coordination of cell fate decisions.

Great endeavors in the late 90s utilized systematic mutagenesis to understand the functional nature of pRb phosphorylation (Barrientes et al., 2000; Brown et al., 1999; Knudsen and Wang, 1997, 1996; Knudsen et al., 1999), pointing to a combinatorial role in pRb regulation (Munro et al., 2012; Rubin, 2013). But this notion has been challenged in recent years based on a report where pRb was found in only three states in cellular lines: un-phosphorylated, hyperphosphorylated and mono-phosphorylated (Narasimha et al., 2014). A more recent report (Sanidas et al., 2019), recapitulated the concept of the phosphorylation code, but focused on mono-phosphorylated isoforms. In that study, the authors found that more than one third of the pRb interactome (434 proteins) bind neither un-phosphorylated nor any of the 14 monophosphorylated variants, and assumed that this portion of the interactions correspond to hyperphosphorylated pRb (Sanidas et al., 2019). Since the phenotypes of our fully phospho-mimetic variant suggest that hyper-phosphorylated RBR is mostly inactive, we believe more evidence is needed before the possibility of intermediate phosphorylation states of RB proteins is rejected. In particular, investigation of pRb phosphorylation states in the whole-organism context remains a future challenge.

To our knowledge, this is the first systematic study of RBR phospho-variants in a full multicellular organismal context. We did not determine the existence of intermediate phosphorylation RBR isoforms, but our “artificial” phospho-variants recapitulated functional outcomes of RBR, implying its potential to “interpret” a combinatorial phosphorylation code modulated by additive effects. We are aware that not all putative phospho-sites have been confirmed *in vivo,* but all phospho-defective variants showed at least a mild and additive effect, implying that at least the majority of phospho-sites are functional. Two shortcomings in our approach are the potential effect of residual RBR in the amiGO genetic background, and the static nature of the phosphorylation substitutions –contrasting with the dynamics entailed by phosphorylation-dependent regulation of a multifunctional protein, achieved by concerted action of CDKs and phosphatases. The former issue could be addressed by postembryonic gene editing to generate a postembryonic rbr null background; the latter would require thorough characterization of the endogenous phosphorylation isoforms of RBR within its diverse spatio-temporal contexts. In this regard, singlecell and single-molecule approaches promise an exciting future for RB protein biology.

Our results support a combinatorial RBR-phosphorylation code in plants and suggest that distinct CYC-CDKs complexes target RBR phospho-sites with distinct affinities as is the case for CYCDs on pRb (Paternot et al., 2006). Accordingly, residues T406 and S911 are preferentially phosphorylated by CYCA3;4, whose overexpression results in RBR-associated phenotypes (Willems et al., 2020); and CYCD6;1, a developmental and stress-responding gene (Bertolotti et al., 2020; Cruz-Ramírez et al., 2012; Zhou et al., 2019), drives the kinase activity of CDKB1 to the Pocket domain (Cruz Ramirez 2012). The distinct expression patterns of CYC genes (Collins et al., 2012; Menges et al., 2005), the intricate regulation of CDK activity (Sanz et al., 2011), and their substrate-specificity, all suggest a complex mechanism to orchestrate RBR multiple functions during the plant life cycle, posing new challenges to our understanding of RBR regulatory networks.

On the same track, RBR-protein interactions mediated by the B-pocket constitute a major component of phosphorylation-mediated functions of RBR (Fig 6), pointing out to proteins containing an LXCXE motif. Noteworthy, RB interactions with LXCXE-containing proteins, both in plants and animals, play prominent roles in sustaining differentiation and growth arrest decisions (Chen and Wang, 2000; Cruz-Ramírez et al., 2012; Matos et al., 2014), and to withstand stressful growth conditions (Bourgo et al., 2011; Collins et al., 2015; Cruz-Ramírez et al., 2013; Zhou et al., 2019). The significant rescue of all phenotypes associated with highly substituted phospho-defective variants by the NF mutation reveals the vast importance of LXCXE protein interactions, but at the same time, it points to an LXCXE-independent component in the phosphorylation-dependent functions of RBR. We showed that RBR interactions with the LXCXE-containing proteins of the TCX5/6/7 clade and with the E2FC proteins, both members of the DREAM complex, are regulated by phosphorylation as they interacted notably stronger with phospho-defective variants than with phospho-mimetics. As the NF substitution conferred stronger suppression of the fully phospho-defective RBR than the *tcx5/6* genetic background, we conclude that several but not all phosphorylation-regulated roles of RBR are intimately linked to the DREAM complex.

Our Y2H analysis suggests new features of RBR previously unknown. First, several interactors are linked to stress and/or environmental responses, like DREB2D to dehydration, high salinity, and heat-stress (Liu et al., 1998)(Chen et al., 2010; Nakashima et al., 2000); GBF4 to cold and dehydration (Lu et al., 1996; Menkens and Cashmore, 1994); NAC090 to reactive oxygen species, salicylic acid responses and sound vibrations that elicit defence hormones (Kim et al., 2018)(Ghosh et al., 2016), the latter being a proposed mechanism to perceive herbivore chewing (Appel and Cocroft, 2014); NAC044 to DNA damage and heat-stress (Takahashi et al., 2019); POB1 to light, vernalization, and susceptibility to pathogens (Christians et al., 2012) (Hu et al., 2014) (Pogoda et al., 2020); and ARIA to the stress hormone ABA (Kim et al., 2004). A subset of these, bind RBR independently of its phosphorylation state, suggesting that hyperphosphorylation does not inactivate RBR completely. Second, the NF mutation disrupted the interaction with TCP3, but only NAC044, and TCX6/7 interact with RBR in a LXCXE-dependent manner (Lang et al., 2021), figS5 S6C); moreover, TCP3 and TCX6/7 interactions with RBR are weakened by the phospho-mimetic mutations. Therefore, phosphorylation may not necessarily block the LXCXE-binding site in RBR as previously thought (Gutzat et al., 2012), and not all proteins binding to this site actually contain an LXCXE motif. And third, RBR-ARIA interaction seems controlled by local rather than global changes in the vicinity of phospho-sites within the N-domain, suggesting a novel mechanism of PPI-regulation by RBR phosphorylation.

Altogether, we have taken first steps to “decipher” the RBR phosphorylation code. Our biochemical and phenotypic analysis suggests that the integrative functions of RBR (Harashima and Sugimoto, 2016) seem to rely on both phosphorylation-regulated and phosphorylation-independent interactions with nuclear proteins. Combinatorial phosphorylation of RBR is essential for developmental processes like (stem) cell division, cell death and differentiation, but seemingly unimportant for binding to several stress-related proteins. Finally, our RBR phospho-variants collections and combinable phospho-modules are a valuable resource for future research. On the one hand, characterization of *in vivo*-phosphorylated sites under environmentally varying conditions may guide the choices to expand the collection. On the other hand, using the collection for cell-type specific, high-throughput experiments and studies on CYC-CDK specificities could help to understand RBR networks throughout development and stress responses. Since RBR is a multifunctional growth regulatory protein, implementation of RBR phosphorylation codes in crops could help to face future food security challenges.

## Materials and methods

### Accession numbers

RETINOBLASTOMA-RELATED (RBR), AT3G12280; G-BOX BINDING FACTOR 4 (GBF4), AT1G03970; E2FC, AT1G47870; Teosinte branched1 Cycloidea1 and PCNA factor 3 (TCP3), AT1G53230; DRE-BINDING PROTEIN 2D (DREB2D), AT1G75490; Tesmin/TSO1-like CXC domain-containing protein 6 (TCX6), AT2G20110; TSL-KINASE INTERACTING PROTEIN 1 (TKI1), AT2G36960; NAC DOMAIN CONTAINING PROTEIN 44 (NAC044), AT3G01600; GROWTH REGULATING FACTOR 5 (GRF5), AT3G13960; POZ/BTB CONTAINING-PROTEIN 1 (POB1), AT3G61600; HD-like, AT4G03250; ARM REPEAT PROTEIN INTERACTING WITH ABF2 (ARIA), AT5G19330; NAC DOMAIN CONTAINING PROTEIN 90 (NAC090), AT5G22380; Tesmin/TSO1-like CXC domain-containing protein 7 (TCX7), AT5G25790; XYLEM NAC DOMAIN 1 (XND1), AT5G64530.

### Plant Material and growth conditions

*Arabidopsis thaliana* ecotype Col-0 was used as wild-type control. Unless otherwise noticed, amiGO-RBR (Cruz-Ramírez et al., 2013) was used as genetic background for transgenic plants. Seeds were fume-sterilized in a sealed container with 100 ml bleach and 3 ml of 37% hydrochloric acid for 3–5 h; then suspended in 0.1% agarose, stratified for 2 d (4 d for arrested embryos) at 4°C in darkness, plated on 0.5x Murashige and Skoog (MS) plus vitamins, 1% sucrose, 0.5g/l 2- (N-morpholino) ethanesulfonic acid (MES) at pH 5.8, and 0.8% plant agar, and grown vertically for 6 d (4 d for arrested embryos) at 22°C with a 16h light/8h dark cycle. For cytological analysis of gametophytes, seedlings were transplanted to soil and grown until reproductive stage.

### RBR phospho-variants plasmid construction and Plant transformation and transgenic selection

The Golden Gate modular cloning (MoClo) system for plants (Engler et al., 2014) was used to generate all phospho-variants. A detailed description of phospho-variants cloning is offered in supplemental methods. Level 2 constructs were transformed in homozygous amiGO plants by flower dip method. Primary tansformants were selected for fluorescent red seed coats under a fluorescence microscope; at least 16 primary transformants were visualized with confocal laser scanning microscopy (CLSM) and selected for the presence of nuclear SCFP3A signal. At least two independent lines were taken to homozygous T3 generation for phenotyping.

### Microscopy

A 10 *μ*g/mL Propidium Iodide (PI) staining solution was used for whole-mount visualization of live roots with CLSM using a Zeiss LSM 710 system as described in (Zhou et al., 2019). For arrested embryos, seed coats were removed as described in (Lee and Lopez-Molina, 2013); modified pseudo-Schiff PI (mPS-PI) staining of roots and embryos was performed as described in (Zhou et al., 2019). Images were taken with ZEN 2012 software (Zeiss) and processed with ImageJ software, using the Stitching Plug-in for multiple images. Brightness and contrast of the final figures was enhanced to the exact same values except for figure S3B,C and S6E where CLSM and brightness/contrast parameters were maximized due to the very weak SCFP3A signal. DIC images of gametophytes were obtained with a Nomarsky illumination Leica DRM system.

### Phenotypic analysis

At least two independent transgenic lines for each genotype were analysed in at least two independent occasions. For the three phenotypes analysed (meristem size, stem cell proliferation and cell death), independent replicates of each line were compared among them; and independent lines of the same genotype we compared among them, with no significant difference. Each line was compared with the Col-0 control, obtaining similar results for each line of the same genotype. Only one line is presented for each genotype. Quantification of the root meristem size was done by imaging the median longitudinal section of the root tip and averaging the number of cortex cells from the QC to the first rapidly elongating cell in the two visible cortex layers; and by measuring and averaging the distance spanned by these cells. Cell death was quantified as the percentage of root tips presenting dead cells as visualized with PI and scanning throughout the Z-axis. Stem cells were visualized by mPS-PI staining and imaging the median longitudinal section of the root tip. Statistical analysis was performed by a One-way ANOVA followed by Dunnett’s multiple comparisons test, or Chi-square followed by Fisher’s exact test using GraphPad Prism version 5.0.0 for Windows, GraphPad Software, San Diego, California USA www.graphpad.com. Histological analysis of female gametophytes was performed with one line per genotype as described previously (Demesa-Arévalo and Vielle-Calzada, 2013).

### Cloning of RBR phospho-variants

All primers used for cloning, carrying relevant restriction sites and 4 bp overhangs were designed using the Primer3 software (Untergasser et al., 2012) and are listed in Table S2. The CDS of Wt RBR was amplified in three fragments, namely N0, AB0, C0, with the primer pairs RBR_n1_WT_F / RBR_n3_WT_R, RBR_AB_F / RBR_AB_R, RBR_C_F / RBR_C_R. Each fragment was cloned in level −1 vector pAGM1311 (generating pAGM1311-RBR_N0, pAGM1311-RBR_AB0, and pAGM1311-RBR_C0). To clone the phospho-defective (N-) and phospho-mimetic (N+) mutant modules, we divided the N fragment it in three sub-fragments (namely n1, n2, n3); fragments n1 and n2 were amplified with primers that introduced the corresponding mutation in the phosphosites T9 and S290: RBR_n1_T9D F / RBR_n1_S290D_R, and RBR_n1_T9A_F / RBR_n1_S290A_R for n1; RBR_n2_S290D_F / RBR_n2_R, and RBR_n2_S290A_F / RBR_n2_R for n2. Mutations in the remaining phospho-sites of module N, were introduced by a synthetic probe, corresponding to n3 fragment, carrying the corresponding mutant codons. The single stranded n3 probes were complemented to double stranded DNA using the n3RvComp primer in a Klenow fragment reaction (Thermo Scientific^™^, EP0421). Phospho-defective and phospho-mimetic fragments n1, n2, n3 were assembled and cloned into pAGM1311 vector to generate N- and N+ modules (pAGM1311-RBR_N- and pAGM1311-RBR_N+). Phosphorylation mutations in the AB and C regions were obtained amplifying the relevant fragments from pre-existing unpublished phosphorylation mutants (generated by serial rounds of directed mutagenesis) with the same primer pairs as for the wild type fragments. The amplicons were cloned into pAGM1311 (pAGM1311-RBR_AB+, pAGM1311-RBR_AB-, pAGM1311-RBR_C+, pAGM1311-RBR_C-). The NT406 mutant modules were obtained by amplifying the corresponding wild type RBR CDS fragment with the primer pairs RBR_n1_WT_F / n3T406A_R, and RBR_n1_WT_F / n3T406E_R, and cloning into pAGM1311 (pAGM1311-RBR_NT406- and pAGM1311-RBR_NT406+). The N849F mutation was introduced in the C0 and C-modules by amplifying two overlapping fragments from the corresponding level −1 modules with the primer pairs RBR_C_F / N849F_R for the first fragment, and N849F_F / RBR_C_R for the second one; both fragments for each C module (wild type and phospho-defective) were assembled and cloned in pAGM1311 vector (pAGM1311-RBRNF_C0 and pAGM1311-RBRNF_C-). All codon changes mentioned above are listed in Table S1.

The combinations of level −1 modules specified in figure S1 were assembled into level 0 vector pAGM1287, creating full length RBR CDS of the corresponding phospho-variant (pAGM1287-RBR_N*AB*C*, where “*” indicates the diverse phospho-modules). For RBR promoter, the intergenic region comprising 1150 bp upstream of the ATG was amplified with primer pair GGpRBR_F / GGpRBR_R, that removed internal BpiI sites, and then cloned into level 0 pICH41295 vector (pICH41294-pRBR). The CDS of the SCFP3A fluorescent protein was amplified with the primer pair GGvYFP_F / GGvYFP_R from an existing clone and sub-cloned into level 0 pAGM1301 vector (pAGM1301-SCFP3A). Each pAGM1287-RBR_N*AB*C* was then combined with pICH41295-pRBR, pAGM1301-SCFP3A, and pICH41421-NosT (from the MoClo toolbox) into level 1 pICH47742 vector (pICH47742-pRBR_RBR_N*AB*C*_SCFP3A_NosT).

Level 1 phospho-variants were cloned into level 2 pAGM4723 vector together with the pICH47732-FAST-R selection marker cassette. All digestion-ligation reactions were performed using 30 fmol of the relevant fragments, plasmids and vector, 1x Green buffer (Thermo Scientific^™^, No.), 1 μM ATP (Thermo Scientific^™^), either 1unit/μL Bsal or Bpil enzymes (Thermo Scientific^™^), T4 DNA ligase (Thermo Scientific^™^) and water to a final volume of 15 μL.

### Plasmid construction for Y2H, transformation, Y2H screenings and confirmation of interactions

We set out to detect RBR protein partners by probing the Y2H library of Arabidopsis transcription factors comprising 1956 nuclear proteins arrayed in 96-well plates (Pruneda-Paz et al., 2014). In total, we used 10 RBR variants as baits: RBR^Wt^, 4 phospho-defective variants, their 4 phosphomimetic counterparts, and RBR^NF^(Fig. S5). pEXP32-RBR-Wt and pEXP32-RBR^NF^ were reported previously (Cruz-Ramírez et al., 2013). The CDS of the eight phospho-variants was amplified from the corresponding level 0 constructs pAGM1287-RBR_N*AB*C* (“*” indicates the diverse phospho-modules listed in Figure S1) with the primer pair cRBR-GWF/ cRBR-GWR and cloned into pDONR221 vector with the Gateway BP clonase II enzyme mix (Invitrogen, 11789020), and the resulting pDONR221-RBR_N*AB*C* phospho-variants entry clones were recombined into pDEST32 by Gateway LR clonase II enzyme mix (Invitrogen, 11791020), resulting in the bait plasmids (pEXP32-RBR_N*AB*C* phospho-variants).

All pEXP32-RBR variants were transformed into yeast strain PJ69-4α and tested for autoactivation as described in (De Folter and Immink, 2011) for at least ten independent transformants; most colonies showed no autoactivation even in selective medium without 3-AT. One colony from each bait with no autoactivation in selective medium supplemented with 0mM 3-AT was inoculated in liquid -L SD-glucose medium and grown O/N. 1mL of the pre-culture was inoculated in 50mL -L SD-glucose medium and grown O/N. In parallel to bait pre-culture, 5 *μ*L of the arrayed Arabidopsis pEXP22-TF library (Pruneda-Paz et al., 2014) was spotted on -W SD-glucose agar plates from PJ69-4A glycerol stock, and grown for 2 days.

A multichannel pipet was used as replicator to transfer the spotted pEXP22-TF library to 96-well plates containing 50 *μ*L of sterile mQ water. 5 *μ*L of the resuspended yeast was spotted on YPD agar plates, letting spots to dry before spotting 5 *μ*L of the pEXP32-RBR variant bait on top and incubating O/N for mating. The yeast was then transferred from YPD to 96-well plates containing 50 *μ*L of sterile mQ water, resuspended, and 5 *μ*L spotted on -LW SD-glucose agar plates and incubated for 3 days to select for the presence of both bait and pray plasmids. The transfer procedure was repeated from the -LW SD-glucose to fresh -LW SD-glucose and −LWH + 1 mM 3-AT SD-glucose agar plates, and incubated for 4-5 days. Selection of positive interaction was based on at least 3 colonies per spot. From the 10 library screenings, we identified 28 interactors in total. To confirm the identity of the positive interactors, plasmid was extracted from the yeast, transformed in chemically competent E. coli DH5-α, selected in LB + ampicillin and mini-prepped again for sequencing.

To assess the interaction patterns, the bait plasmids (pEXP32-RBR variants) were transformed into yeast strain PJ69-4A as described in (De Folter and Immink, 2011). No autoactivating colonies carrying each of the baits were made competent and transformed with the purified and sequenced pEXP22-TF obtained from the screenings. The transformation was adapted from (De Folter and Immink, 2011) to be done in 96 deep well plates instead of 1.5 mL tubes. Transformed yeast was resuspended in 150 *μ*L of sterile mQ water and spotted in triplicate on -LW SD-glucose agar plates and incubated for 3 days. The yeast was then transferred to 96-well plates containing 50 *μ*L of sterile mQ water, resuspended, and 5 *μ*L spotted onto fresh -LW SD-glucose, –LWH + 1.5 mM 3-AT SD-glucose, and -LWHA SD-glucose agar plates, and incubated for 5 days. Selection of positive interaction was based on at least 3 colonies per spot.

For pEXP22-TCX6^gcg1^ and pEXP22-TCX6^gcg2^, we first generated the corresponding entry clones by amplifying the attB-flanked TCX6 CDS in a two-fragments overlapping PCR to mutagenize the LXCXE and LXCXE-like motifs with the primer pairs TCX6c-F/ TCX6gcg1-R and TCX6gcg1-F /TCX6c-R for TCX6^gcg1^; and TCX6c-F/TCX6gcg2-R and TCX6gcg2-F /TCX6c-R for TCX6^gcg2^, followed by BP-II clonase (Invitrogen) recombination reaction into pDONR-221. Entry clones were recombined into pDEST22 destination vector with the LR-II clonase (Invitrogen). Small scale Y2H assays was performed by co-transforming bait and pray plasmids as described in (De Folter and Immink, 2011), testing 12 independent co-transformations per interaction. All yeast incubations described above were at 30 °C. Plates were imaged at with a table top flatbed scanner (EPSON Expression 11,000 XL).

## Acknowledgements

We thank Prof. Jim Murray (Cardiff University) for his valuable suggestions on an early version of this manuscript; Dr. Xin-Jian He (National Institute of Biological Sciences, China) for sharing their *tcx5/tcx6* mutant line; Dr. Sara Díaz-Triviño (Wageningen University) for sharing the unpublished RBR^NF^-YFP line, Dr. Yessica Alina Rodríguez-Rosales (Radboud UMC) for her support with statistical analysis and R plots, Jonathan Matthew Samson and Dr. Renze Heidstra for technical support. JZZ was funded by Consejo Nacional de Ciencia y Tecnología (CONACyT, Mexico, 383871).

## SUPPLEMENTARY FIGURE LEGENDS

**Figure S1.**
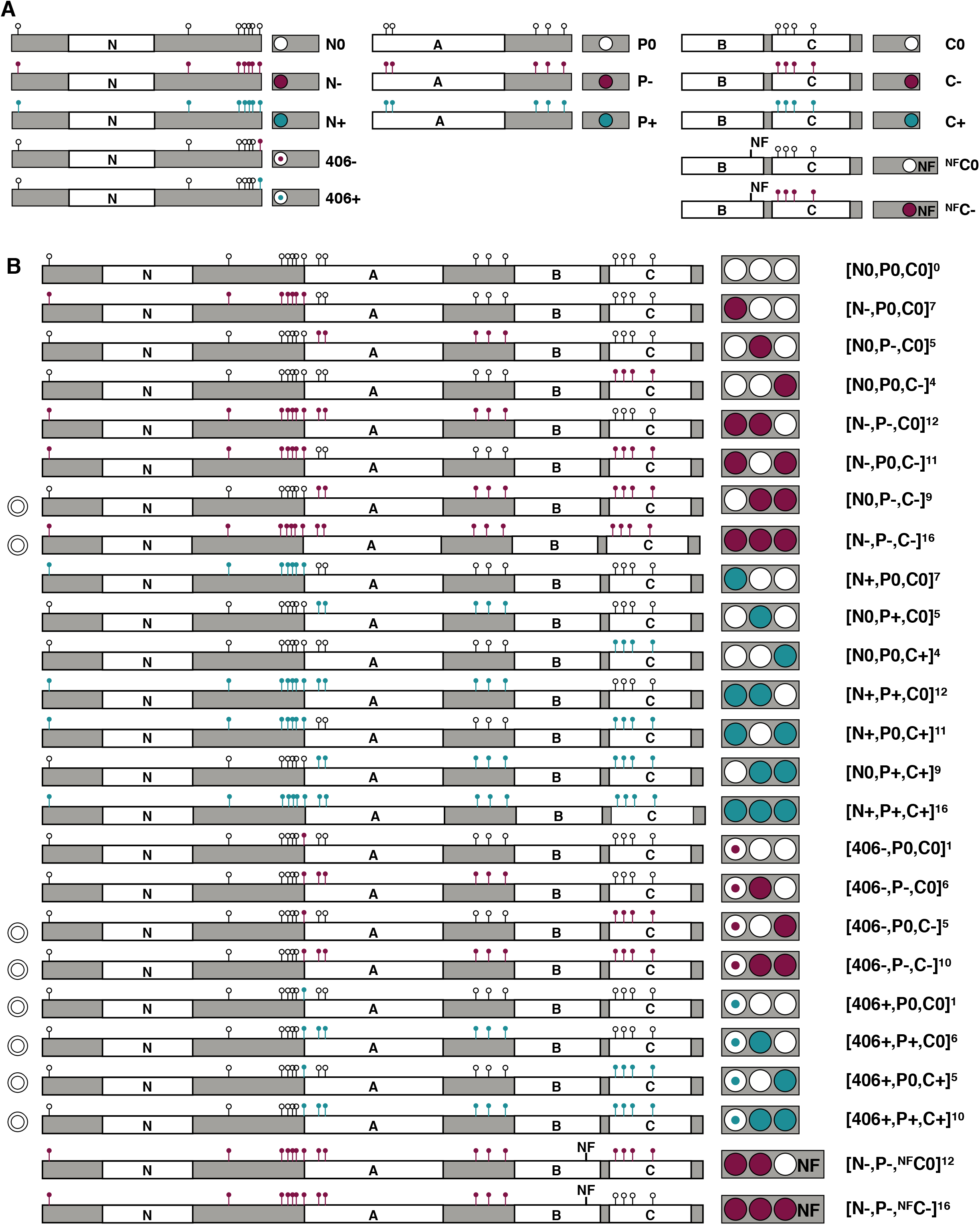
List of modules and RBR-phosphovariants. A) Schematic list of all modules cloned in Level −1 (vector pAGM1311) of the GoldenGate MoClo system. Text nomenclature and coloured circles and their relative position within the gray background box indicate the phosphorylation state and position of each module within the full length CDS of phosphovariants according to Fig 1. Note that in modules “406” only the Thr406 residue is mutated, and in modules “^NF^C” the Asn849 within the LXCXE binding cleft of the B-pocket sub-domain is mutated to Phe. B) Schematic list of all phospho-variants generated. All variants listed exist as Level 0 (vector pAGM1287), Level 1 (vector pICH47742; with RBR promotor, SCFP3A CDS and NOS terminator), and Level 2 (vector pAGM4723; with FAST-R selection cassette in position 1) constructs, and as transgenic seed, except for those marked with the symbol ◎ on the left-most column, which were either not viable or not transformed and thus, only the plasmids are available. Note that [406-,P0,C-]^5^ is marked with double circle because reduced fertility hindered propagation and all seed was used for the experiments reported in Fig 5.

**Figure S2.**
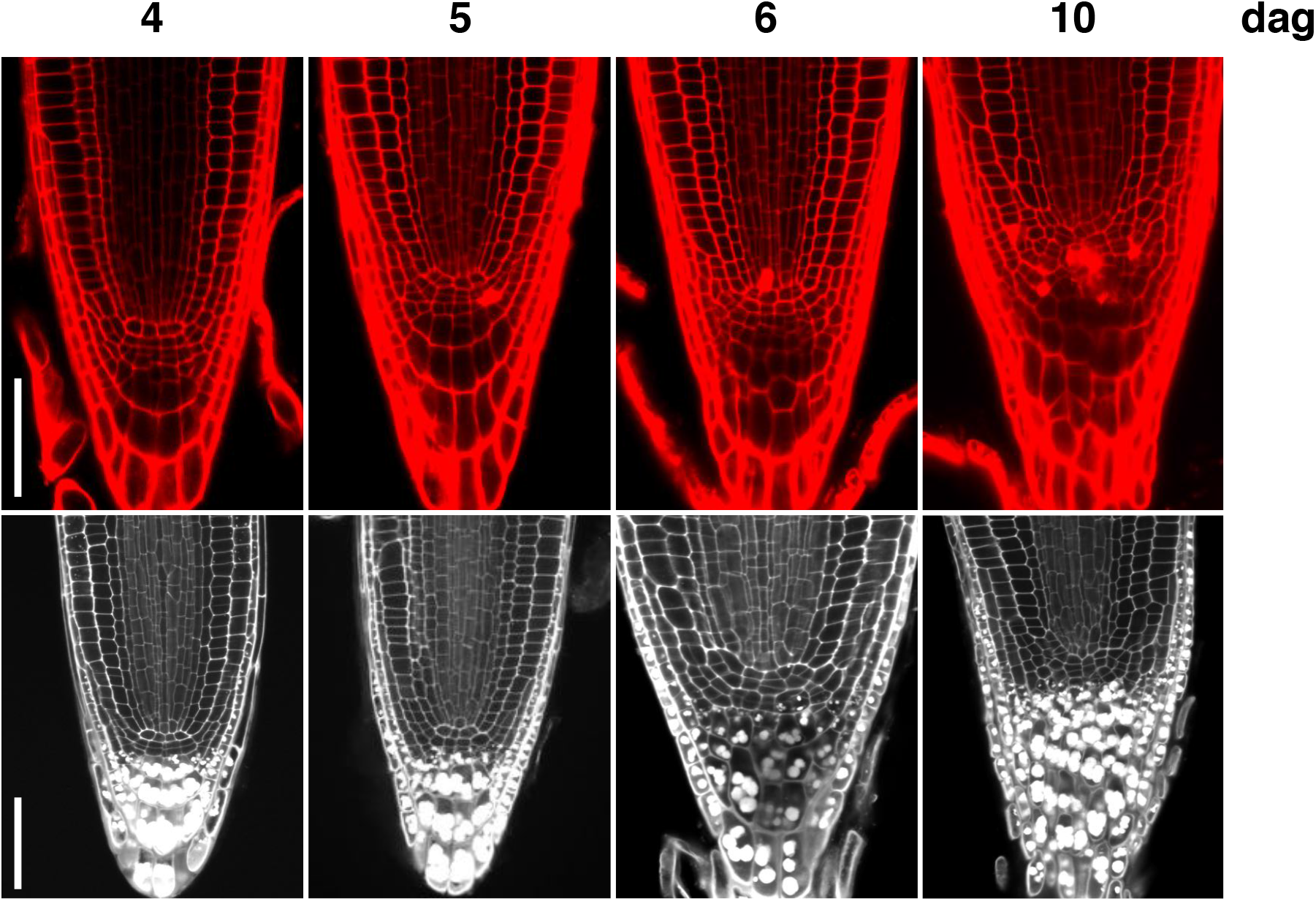
amiGO-RBR phenotype penetrance. Confocal images of amiGO-RBR root tips by 4,5, 6 and 10 days after germination (dag). Top panels, mPS-PI staining; bottom panels, PI staining. Note that by 5 das we detected roots with very weak or no amigo-associated phenotypes. By 6das, cell death and SCN extra divisions were evident in all roots. Scale bars, 50 μM.

**Figure S3.**
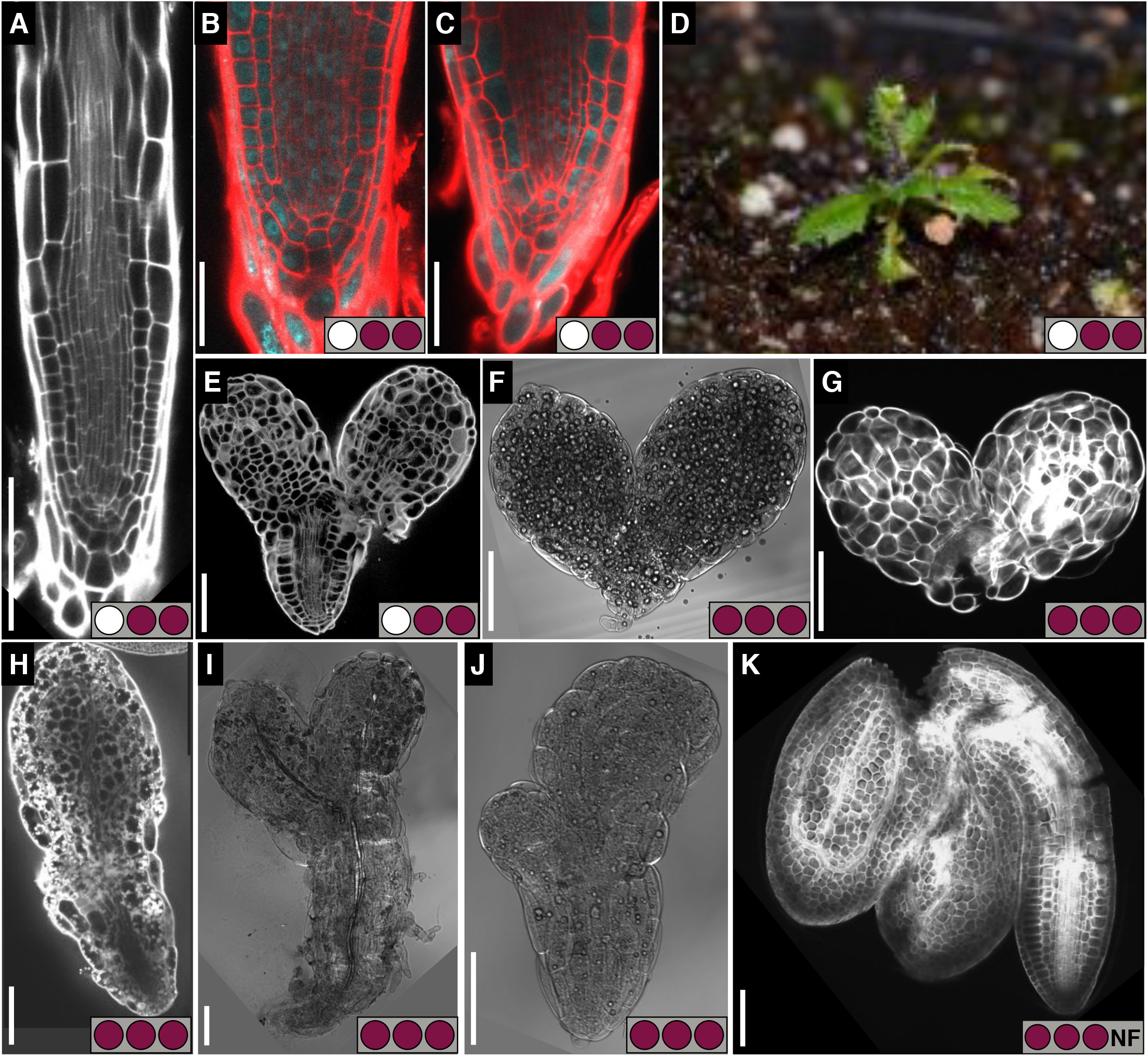
Primary transformants of lethal phospho-defective RBR variants. A-C,E-K) Confocal images of PI stained root tips A-C), mPS-PI stained E,G,H,K), and transmitted light images F,I,J) of embryos from non-germinated seeds 4 das stratified for 4 days. D) 3 week old seedling. Genotypes: [N0,P-,C-]^9^ A-E), [N-,P-,C-]^16^ F-J), [N-,P-^NF^,C-]^16^ K); Genetic background: Col-0 A,B,G,K), amiGO C-F,H,-J). Max power and gain for CLSM settings and image brightness and contrast in B,C) were set to in order to visualize sCFP signal from [N0,P-,C-]^9^ expression. Scale bars, 100 μM in (A,E-K), 50 μM in (B,C).

**Figure S4.**
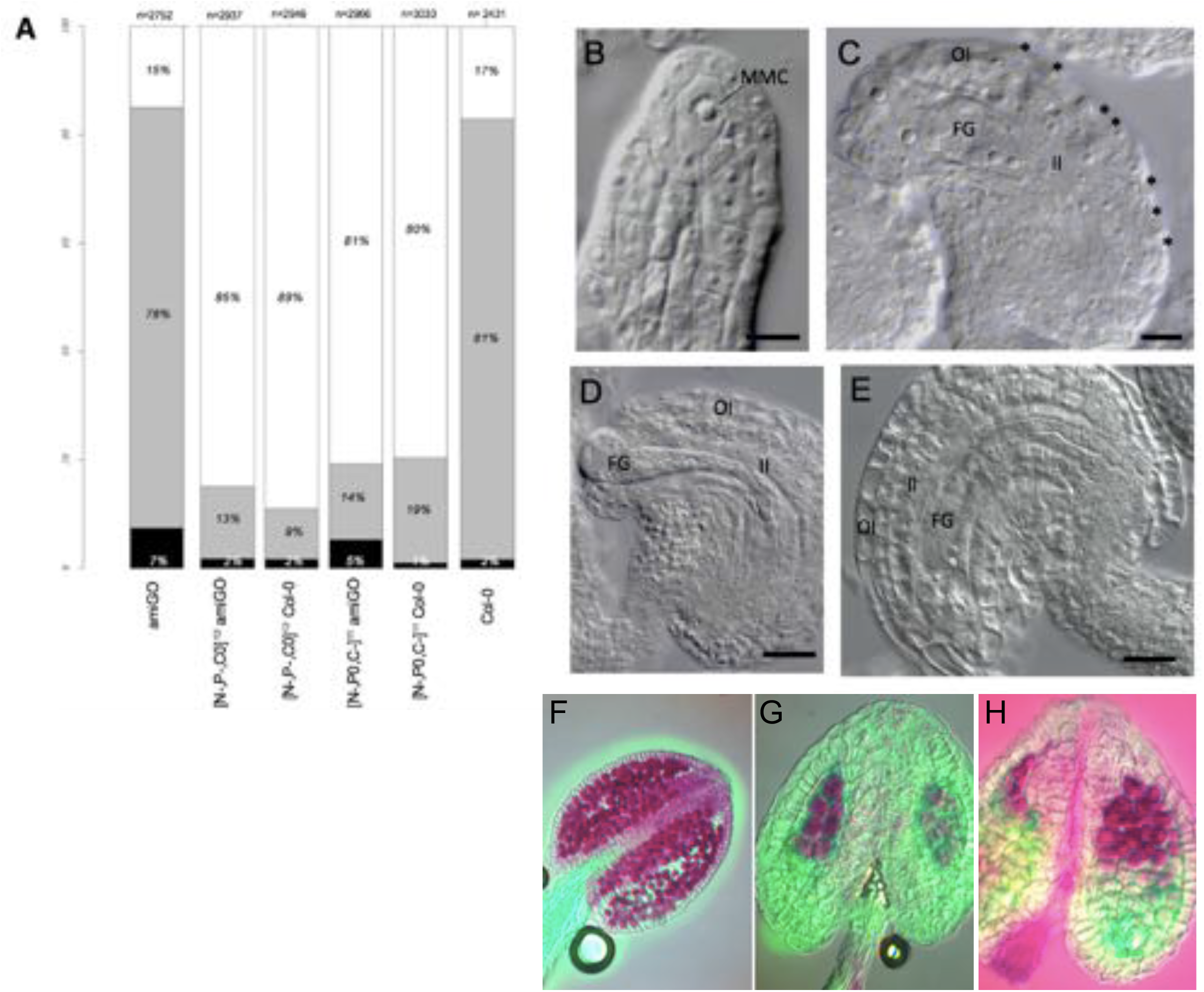
Fertility and embryogenesis are compromised in the highly substituted RBR phospho-defective variants [N-,P-,C0]^12^ and [N-,P0,C-]^11^. A) Sterility analysis quantified as percentage of non-fertilized ovules, aborted seed and mature seed. N denotes total number of scored ovules and seed. B-K) DIC images of ovule development of [N-,P-,C0]^12^ B-F) and [N-,P0,C-]^11^ genotypes. Ovule primordia with one B,G) or more C,H) MMC. D-F,I-K) Incomplete integument development results in abnormally exposed embryo sac. L-N) Alexander staining of Col-0 L), [N-,P-,C0]^12^ M), and [N-,P0,C-]^11^ N) anthers showing viable pollen grains in fuchsia.

**Figure S5.**
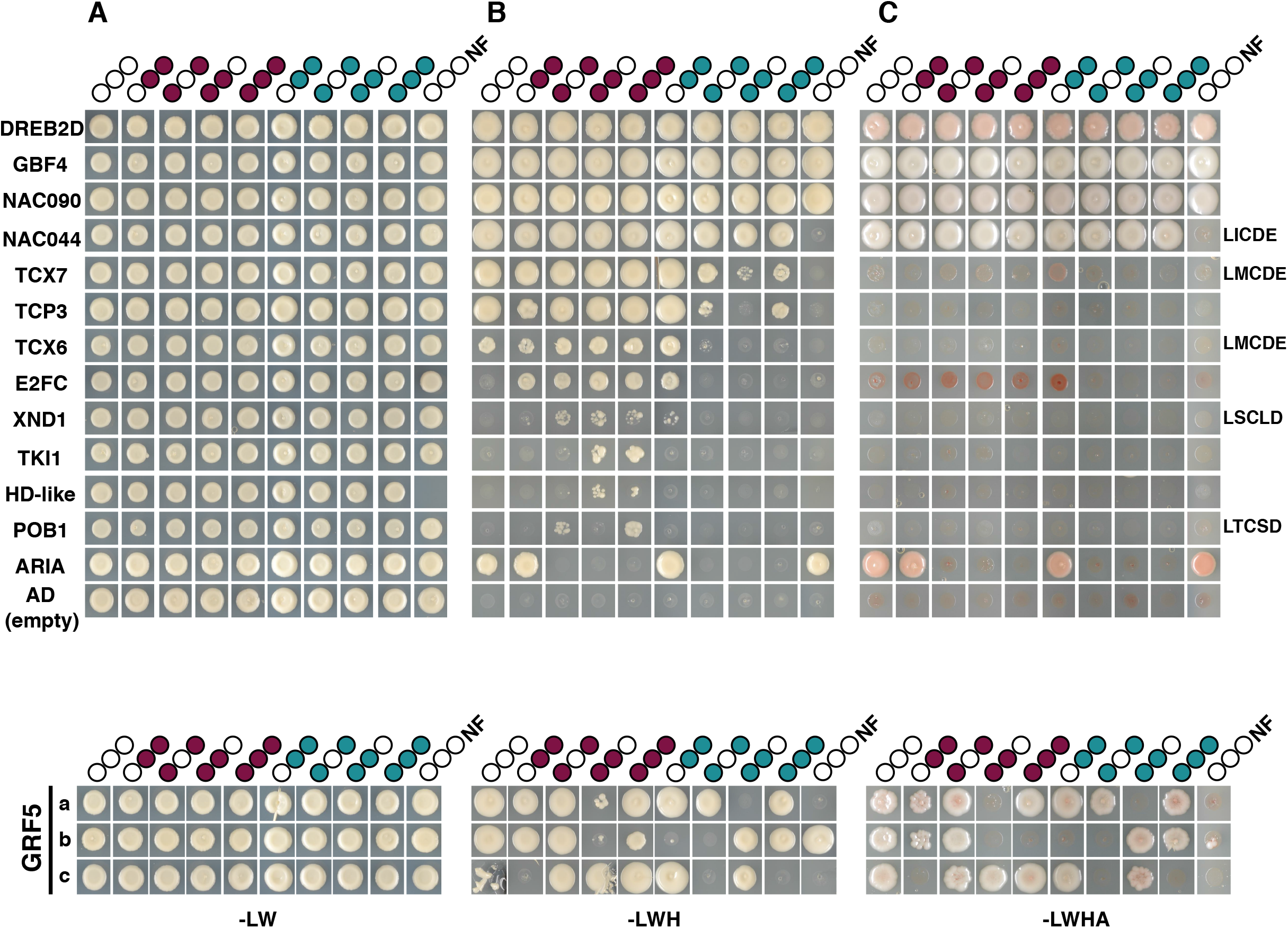
RBR protein-protein interactions with transcriptional regulators are differentially regulated by phosphorylation as shown by Y2H screenings of the Arabidopsis transcription factors library. A) Y2H analysis of co-transformed pEXP32-RBR variants and pEXP22-TFs dropped on SD-LW A), SD-LWH +1.5mM 3AT B) and SD -LWHA C). The identity of each TF fused to the GAL4 activating domain is indicated on the left column; if present, LXCXE and LXCXE-like motifs sequences are indicated in the rightmost column; the total number of interactions per RBR variant is indicated on the lower-most row. Negative control is the empty pDEST22 vector. B) From the Y2H analysis, the three replicates denoted by lowcase letters show that the GRF5 TF interacted strongly but inconsistently to all RBR variants. Cotransformed yeast dropped on SD -LW to select transformants, and on SD –LWH +1.0 mM 3AT and SD -LWHA to select interactions.

**Figure S6.**
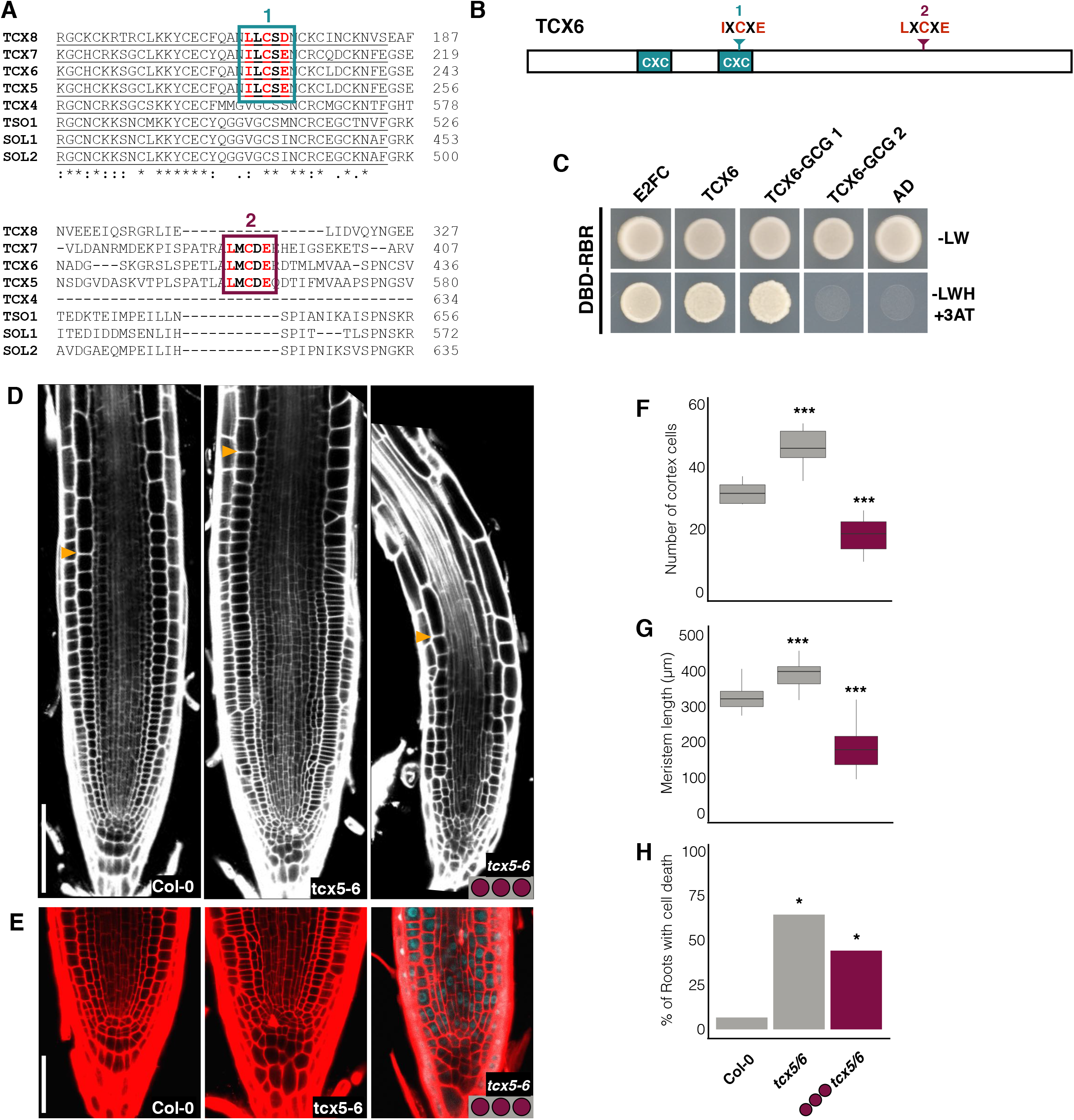
Phosphorylation-regulated functions of RBR are largely mediated by the interaction with members of the DREAM complex. A) Clustal omega multiple sequence alignment fragments of Arabidopsis TCX proteins showing LXCXE-like motifs (red) within the conserved CXC domain (underlined) and LXCXE motifs (in green) within a less conserved region; asterisks and dots indicate identical and similar residues, respectively. B) Schematic representation of TCX6 protein organization as predicted by Pfam server (https://pfam.xfam.org/), showing the relative positions of the cysteine-rich domains (cxc) and LXCXE and LXCXE-like motifs. C) Yeast two-hybrid analysis showing that RBR interacts with TCX6 and a TCX mutated on the LXCXE-like motif ‘1’ within the CXC domain, but not with TCX6 mutated on the canonical LXCXE motif ‘2’. E2FC is positive control, and empty pDEST22 vector is negative control. Co-transformed yeast dropped on SD −LW to select transformants, and on SD -LWH +1.0 mM 3AT to select interactions. D,E) Representative confocal images of PI-stained root tips of the indicated genotypes; yellow arrowheads mark the end of the meristem proliferation zone. The CLSM settings for detecting SCFP3A in the *tcx5/6*;[N-,P-,C-]^16^ were identical than those for all other phospho-variants, but brightness and contrast were enhanced to visualize the nuclear signal due to the low fluorescence intensity. F,G) Box plots from D) of meristem proliferation and size quantified as the number of cortex cells F) and length G) from the QC to the first rapidly elongating cortex cell. H) Bar graph from E) showing the percentage of root tips with dead cells. Data from one biological replicate presented as median and interquartile range F,G), or as means H); n>15. Wilcoxon test against Col-0, ***p < 0.001 in F,G), Chi square, *p < 0.05 in H). Scale bars, 100 μM in D), 50 μM in E).

**Supplementary Table S1.**
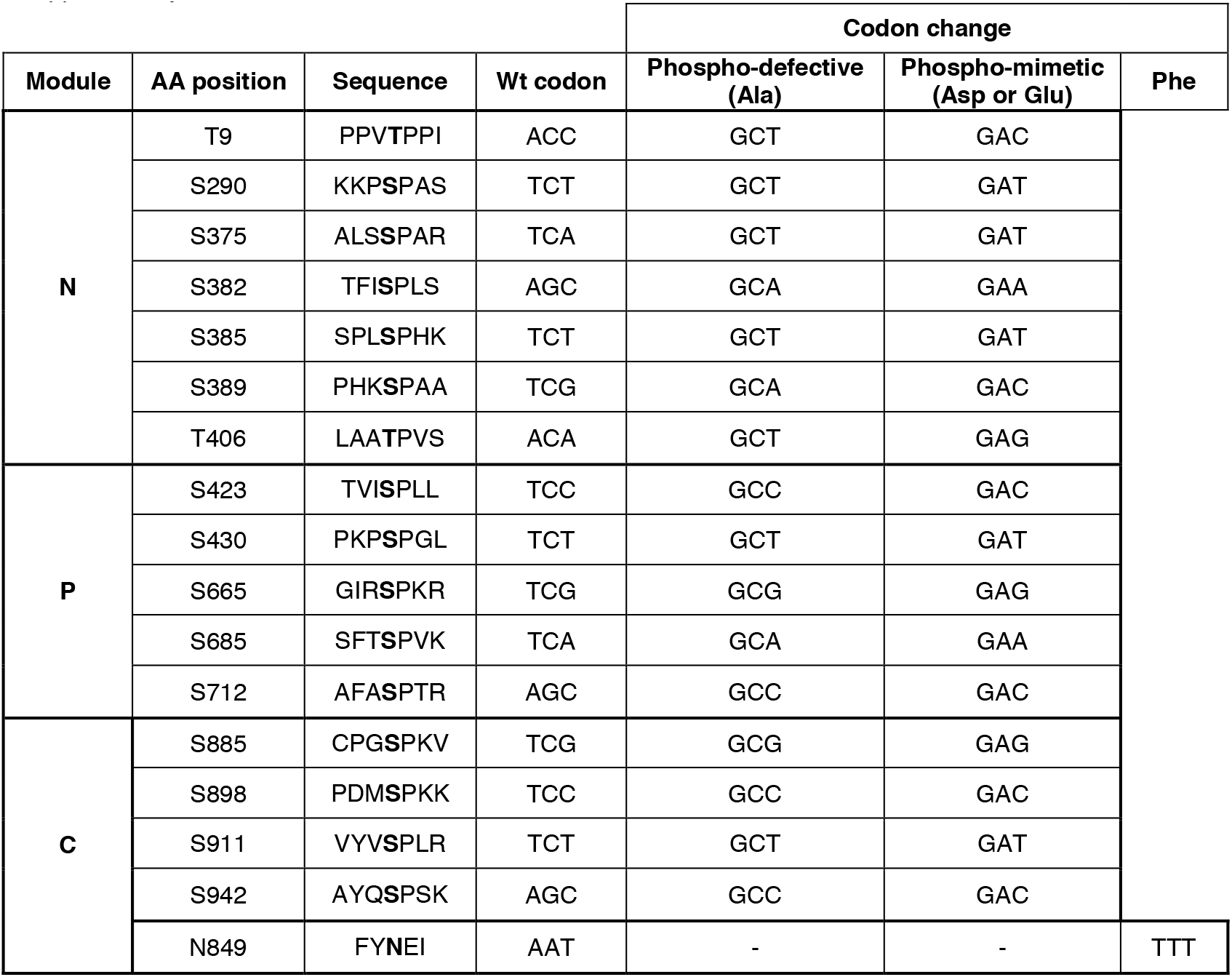
Codon changes in RBR amino acid substitutions.

**Supplementary table S2.**
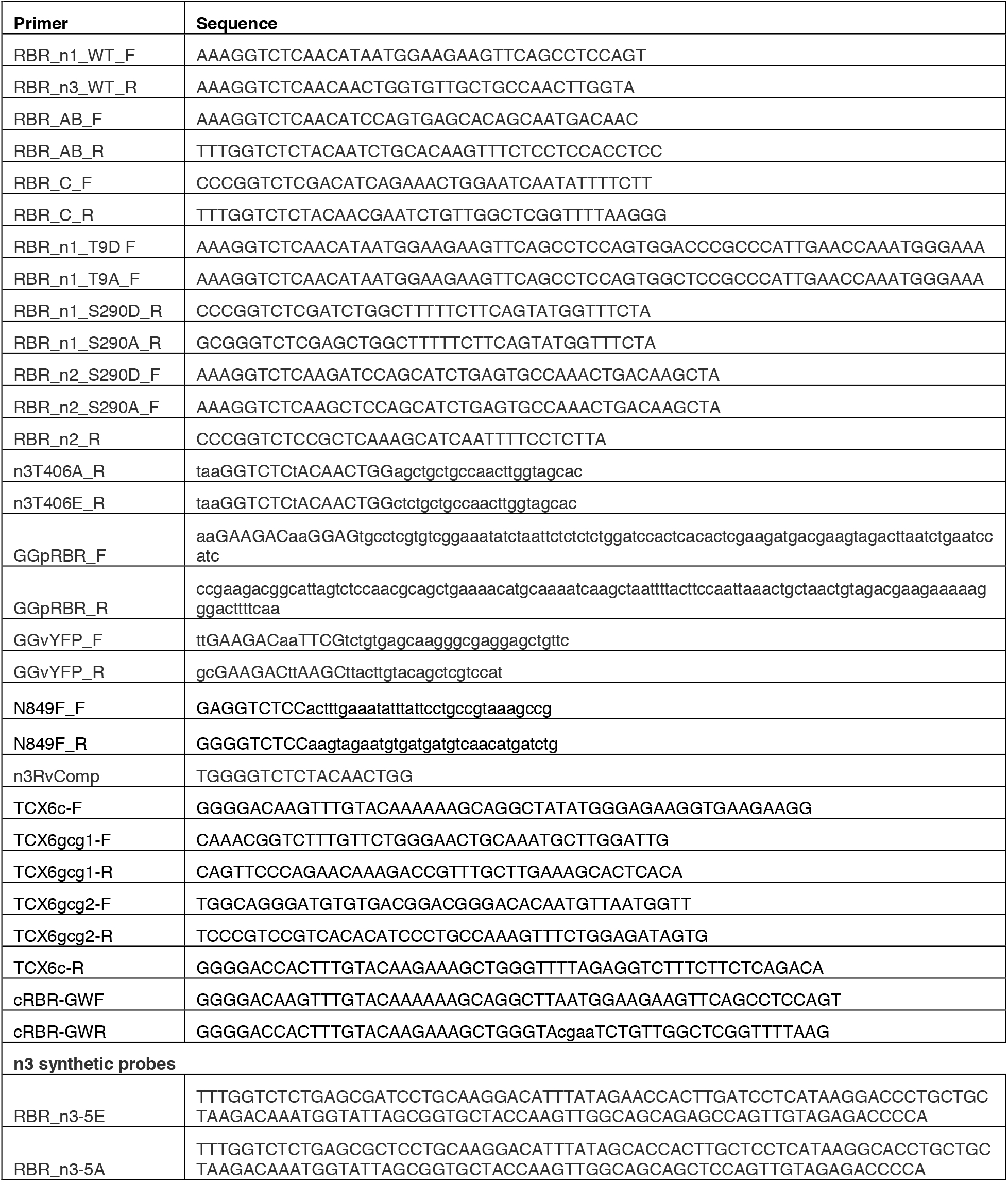
List of primers

## Notes

### Competing Interest Statement

The authors have declared no competing interest.

